# The Folding of Germ Granule mRNAs Controls Intermolecular Base Pairing in Germ Granules and Maintains Normal Fly Development

**DOI:** 10.1101/2024.05.31.596852

**Authors:** Siran Tian, Hung Nguyen, Ziqing Ye, Silvi Rouskin, D. Thirumalai, Tatjana Trcek

## Abstract

*Drosophila* germ granules enrich mRNAs critical for fly development. Within germ granules, mRNAs form multi-transcript clusters marked by increased mRNA concentration, creating an elevated potential for intermolecular base pairing. However, the type and abundance of intermolecular base pairing in mRNA clusters is poorly characterized. Using single-molecule super-resolution microscopy, chemical probing for base accessibility, phase separation assays, and simulations, we demonstrated that mRNAs remain well-folded upon localization to germ granules. While most base pairing is intramolecular, mRNAs still display the ability for intermolecular base pairing, facilitating clustering without high sequence complementarity or significant melting of secondary structure. This base pairing among mRNAs is driven by scattered and discontinuous stretches of bases appearing on the surface of folded RNAs, providing multivalency to clustering but exhibits low probability for sustained interactions. Notably, engineered germ granule mRNAs with exposed GC-rich complementary sequences (CSs) presented within stable stem loops induce sustained base pairing *in vitro* and enhanced intermolecular interactions *in vivo.* However, the presence of these stem loops alone disrupts fly development, and the addition of GC-rich CSs exacerbates this phenotype. Although germ granule mRNAs contain numerous GC-rich CSs capable of stable intermolecular base pairing, they are primarily embedded by RNA folding. This study emphasizes the role of RNA folding in controlling the type and abundance of intermolecular base pairing, thereby preserving the functional integrity of mRNAs within the germ granules.

## Introduction

Membraneless biomolecular condensates enrich proteins and RNAs, which spatiotemporally regulate post-transcriptional gene regulation (*1, 2*). These condensates have been observed across species (*3–8*) and often appear structured, with their proteins (*9–15*) and RNAs (*3, 16–20*) condensing into clusters consisting of multiple macromolecules, *in vivo* and *in vitro*.

Protein conformation and protein:protein interactions within proteins clusters have been extensively studied (*21–24*). Depending on the species, some proteins, like Poly(A)-binding protein (Pab1) and globular proteins, partially or completely unfold during condensation (*21, 24*). Others, such as Fused in Sarcoma (FUS) and androgen receptor activation domain (AR AD), transition to a more compact and folded state (*22, 25*). Regardless of their conformations, these clusters require multivalent interactions driven by charge-charge, cation-π, π-π, dipole-dipole and hydrophobic interactions, facilitated by diverse amino acid sequence compositions (*23, 26, 27*).

However, our understanding of RNA:RNA interactions and RNA conformations within RNA clusters remains limited and is mainly derived from *in vitro* studies. In these studies, *in vitro* transcribed RNAs self-assemble into microscopically visible clusters with the addition of salts and crowding reagents. This clustering occurs in the absence of other cellular components (*16, 19, 20, 28, 29*) demonstrating the potential of RNAs to interact with each other. Similar to clustering of proteins, RNA clustering requires interaction multivalency (*30*) and may involve sequence complementarity. For example, in the filamentous fungus *Ashbya gossypii*, the co-clustering of *BNI1* and *SPA2* mRNAs is facilitated by base pairing of exposed complementary sequences (“zipcodes”) shared between the two transcripts located in five distinct RNA regions (*28*).

To facilitate RNA clustering, intermolecular base pairing may require melting of the RNA structure. Notably, repeat RNAs that cause human neurodegenerative disorders (*31–33*) contain an array of exposed and repeated GC-rich sequences that induce RNA condensation *in vitro* and *in vivo* (*16*). Computer simulations of these RNAs revealed that in the process of condensation, the hairpin structures formed by expanded repeats transition into an unwound state (*34*). This transition enables base pairing among RNAs that increases multivalency of interactions and the formation of an extended network, which restricts RNA mobility within clusters, thereby stabilizing it (*16, 34*). Similarly, guanidine riboswitches (*29*) melt their secondary structures to augment intermolecular base pairing *in silico* or *in vitro*. Finally, antisense non-coding RNAs (ncRNAs) are thought to enrich in stress granules by melting their secondary structures to facilitate intermolecular base pairing with their sense mRNA partners (*20, 30*). Here, the increased length of ncRNAs could result in more extensive complementary base pairing, thereby enhancing multivalent interactions.

*In vivo*, mRNAs critical for development of *Drosophila* enrich in germ granules at the posterior of the developing embryo where they are post-transcriptionally regulated (*35*). These mRNAs self-organize within a homogeneously distributed protein environment of germ granules (*17, 36*) at the posterior of the developing embryo (*35*) forming clusters that contain multiple mRNAs derived from the same gene (*17, 18, 37*). The mRNA concentration within these clusters is remarkably high (5-15µM) (*36*) suggesting dense packing of mRNAs and potential involvement of intermolecular base pairing in their formation.

Interestingly, previous study using chimeric experiments and stochastic super-resolution microscopy (STORM) on germ granule mRNAs failed to detect dependence of RNA clustering on a particular RNA zipcode (*36*). These findings differ from observations of *BNI1* and *SPA2* mRNAs, as well as the *Drosophila oskar* (*osk), bicoid* (*bcd*) mRNAs and *human immunodeficiency virus (HIV)* genomes. Clustering of these RNAs relies on specific, often GC-rich complementary sequences (CSs), that are exposed on stem-loops. This structural arrangement facilitates recognition among mRNAs, leading to stable intermolecular base pairing and RNA clustering (*36, 38–44*). These results highlight the distinctive nature of intermolecular base pairing in *Drosophila* germ granules, setting it apart from the mechanisms observed in *BNI1, SPA2*, *osk, bcd* mRNAs and *HIV* genomes. However, the type and abundance of intermolecular base pairing governing clustering of the *Drosophila* germ granule mRNAs remains poorly understood.

Applying *in vivo*, *in vitro*, and *in silico* experiments, we now demonstrate that contrary to repeat RNAs, germ granule mRNAs remain well-folded upon localization to germ granules, while retaining their capacity for intermolecular base pairing *via* exposed bases appearing on the surface of folded RNA conformations. These interactions occur independently of high sequence complementarity or significant melting of secondary structure and exhibit a low probability for sustained base pairing. This result explains the lack of dependence of RNA clustering on a particular RNA sequence in germ granules we reported previously (*36*). Although appearing more dynamic, these interactions retain interaction multivalence reminiscent of the one driving protein condensation. Finally, engineered germ granule mRNAs with exposed GC-rich CSs presented within stable stem loops induce persistent base pairing *in vitro* and enhanced intermolecular interactions *in vivo* but also disrupt normal fly development. Notably, while the presence of a strong stem loop is the major driver of developmental defects, the exposed GC-rich CSs exacerbate them. Our study reveals that germ granules may employ RNA folding as a mechanism to fend off potential detrimental effects of exposed GC-rich CSs, while facilitating multivalent intermolecular interactions and mRNA clustering. This regulation preserves the normal gene function of germ granule mRNAs and supports fly development.

## Results

### Efficient RNA clustering can occur without extensive sequence-complementarity or significant melting of RNA structure *in vitro*

We first examined the type of intermolecular interactions within the clusters of germ granule mRNAs *in vitro*. To this end, we used the split broccoli system coupled with the chemical reporter DFHBI-1T (*45*). This system involves *top* and *bottom* RNAs, each carrying half of the broccoli aptamer. When these two RNA segments dimerize, they form a broccoli structure detectable by fluorescence emitted by DFHBI-1T intercalated between base-paired *top* and *bottom* RNAs (Figure 1a) (*45*). We reasoned that the observation of a strong broccoli fluorescent signal will report on intermolecular base pairing driven by extensive sequence complementarity as well as reveal unfolding of the secondary RNA structure during clustering.

**Figure 1:**
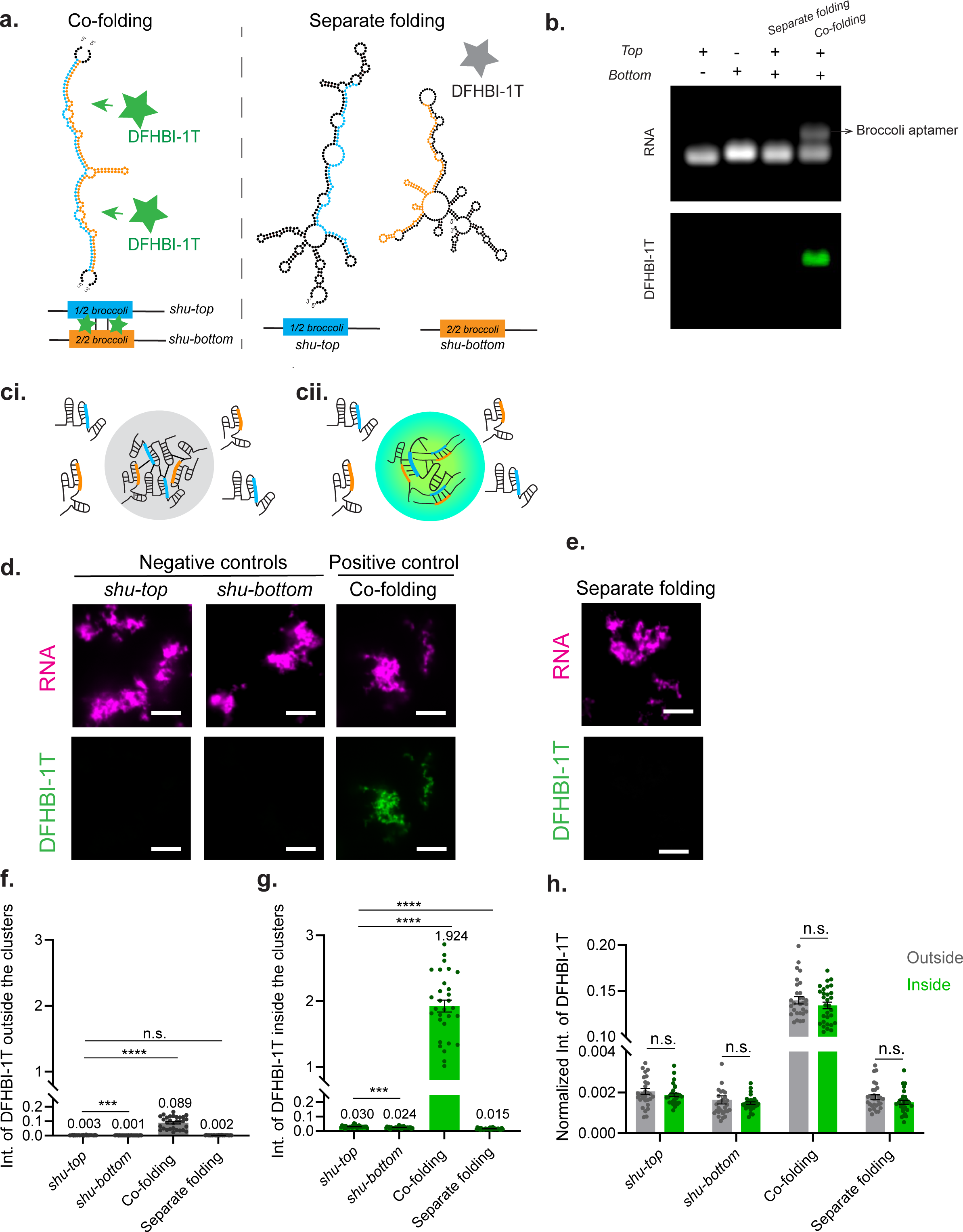
Efficient RNA clustering can occur without extensive sequence-complementarity or significant RNA structural melting *in vitro*. **a.** Schematic and predicted structures of *shu-top* and *shu-bottom* RNAs. *shu-top* and *shu-bottom* RNAs contain split-broccoli sequences *top* (blue) and *bottom* (orange), and their intermolecular base pairing is recognized by DFHBI-1T (see Table S1 for sequences). **b.** *In vitro* base pairing assays for *shu-top* and *shu-bottom* RNAs. Top panel: RNA; bottom panel: DFHBI-1T staining. **c.** Schematics of two distinct models for clustering of *shu-top* and *shu-bottom* RNAs. If RNA clustering occurred without significant structural melting and independently of base pairing driven by extensive sequence complementarity between *shu-top* and *shu-bottom* RNAs, then DFHBI-1T would not emit strong fluorescence inside the cluster (grey) (i). However, if clustering involved melting of the RNA structure and extensive base pairing between the two RNAs, we expect to observe high DFHBI-1T fluorescence inside the clusters (green) (ii). d-e. *In vitro* RNA clusters (magenta) formed of only *shu-top* or *shu*-*bottom* RNAs (negative controls), co-folded (positive control) or separately folded *shu-top* or *shu*-*bottom* RNAs (**e**) in the presence of DFHBI-1T (green). The DFHBI-1T images in **d-e** were normalized using the parameters of the co-folded DFHBI-1T image (see also Figure S1b for images normalized to the DFHBI-1T signal of *shu-top*). Initial RNA concentrations: 3.2 μM for negative controls, 1.6 μM for each RNA in co-folding and separate folding. Scale bars: 7.5 μm. **f-g.** Intensity of DFHBI-1T outside (**f**) or inside (**g**) the microscopic clusters (d, e: magenta). n=30 for all conditions. Data: Mean±SEM. The numbers shown above the bars represent the average intensity values. Although the differences among the negative controls and separate folding were significant, their intensity values were much more similar compared to co-folding, our positive control. n.s.: not significant. ***, ****: p < 0.001 and 0.0001, respectively (unpaired t-test). **h.** Normalized intensity of DFHBI-1T by RNA intensities (Figure S1a) from outside (grey) or inside (green) in RNA clusters per condition. n=30 for all conditions. Data: Mean±SEM. n.s.: not significant (unpaired t-test).

We integrated *top* and *bottom* RNA segments into the *3′UTR* of the *Drosophila shutdown* (*shu*), a germ granule mRNA (*46*), creating *shu-top* and *shu-bottom* (Figure 1a, Table S1). RNAcofold (*47*) predicted the correct broccoli structure when *shu-top* and *shu-bottom* were co-folded, while RNAfold (*47*) predicted that *top* and *bottom* segments base paired intramolecularly when they were folded separately (Figure 1a).

To validate the RNAcofold and RNAfold predictions that *shu-top* and *shu-bottom* form correct broccoli structure detectable by DFHBI-1T, we used non-denaturing RNA gel electrophoresis following established protocols (*39, 41*). After heat-denaturing and co-folding *shu-top* and *shu-bottom* RNAs, we observed a dimerized broccoli RNA accompanied by a strong DFHBI-1T fluorescence on the gel (Figure 1b). In contrast, separately folded and subsequently mixed *shu-top* and *shu-bottom* RNAs did not dimerize and showed no reporter signal, similar to *shu-top* and *shu-bottom* alone, our negative controls (Figure 1b). This absence of fluorescence indicated that even at high RNA concentrations (3.2μM), *shu-top* and *shu-bottom* RNAs did not form an intermolecular broccoli structure when folded separately. In addition, lack of a visible dimer band in samples containing only *shu-top* and *shu-bottom* RNAs, or when the two RNAs were folded separately and afterwards mixed, revealed that *shu* RNA lacked the potential for stable dimerization (Figure 1b).

We then examined if clustering of *shu-top* and *shu-bottom* relies on base pairing among complementary sequences accompanied by unfolding of the RNA structure as observed for repeat RNAs (*16, 34*). To this end, we used fluorescently labeled *shu-top* and *shu-bottom* RNAs and adapted existing RNA phase separation protocols (*20, 28*). We reasoned that if RNA clustering occurred without significant structural melting and independently of base pairing driven by extensive sequence complementarity, we would observe similar DFHBI-1T fluorescence levels inside and outside the *shu-top* and *shu-bottom* clusters (Figure ci). However, if clustering of *shu-top* and *shu-bottom* RNAs involved structural melting and extensive base pairing, we expect to observe a much higher DFHBI-1T fluorescence inside the clusters compared to outside (Figure 1cii). Importantly, we estimated the ratio between DFHBI-1T and *shu-top*/*shu-bottom* RNA dimer of 62.5:1 in our reactions. This ratio ensured a large excess of the fluorescent reporter compared to the dimerized RNA and saturated detection of broccoli structures (see methods (*48*)). Therefore, the intensity of the DFHBI-1T fluorescence should scale with the abundance of broccoli structure.

To establish the baseline for the DFHBI-1T fluorescence inside and outside of RNA clusters, we first measured DFHBI-1T fluorescence in RNA clusters formed by *shu-top* and *shu-bottom* alone (Figure 1d, green signal in images; negative control). Importantly, the raw DFHBI-1T fluorescence outside (Figure 1f) and inside (Figure 1g) of RNA clusters for both RNAs was minimal compared to the positive control (see below), setting the baseline for background fluorescence of DFHBI-1T. DFHBI-1T fluorescence was 10- or 24-fold higher inside the clusters of *shu-top* (0.030±0.002) or *shu-bottom* (0.024±0.001), respectively, compared to the outside (0.003±0.0003 and 0.001±0.0001, respectively). However, after normalizing the fluorescence to their respective RNA intensity inside and outside the clusters (Figure S1a; see methods), we did not observe significant difference in DFHBI-1T fluorescence between inside and outside the clusters for either negative control (Figure 1h). This suggests that the increased DFHBI-1T intensity in RNA clusters formed by the two negative controls was primarily due to the higher RNA concentration inside the clusters, leading to an increased background signal. These results were anticipated given that strong DFHBI-1T fluorescence can only form when *shu-top* and *shu-bottom* dimerize with each other (Figure 1b).

As a positive control, we then phase separated co-folded *shu-top* and *shu-bottom* RNAs. As anticipated, we observed a 30-to 89-fold increase in DFHBI-1T fluorescence outside the clusters, and a 64-to 80-fold increase inside the clusters, compared to *shu-top* or *shu-bottom,* respectively (Figure 1d-g, S1b). For co-folding, the fluorescence inside the clusters was 21-fold higher compared to outside indicating that more dimerized broccoli RNA accumulated within clusters (Figure 1f, g). However, after normalization to the RNA intensity (Figure S1a), we observed no significant difference in DFHBI-1T levels between inside and outside of clusters (Figure 1h). Therefore, the increased RNA concentration within the microscopic clusters did not result in additional broccoli formation when *shu-top* and *shu-bottom* RNAs were co-folded. Importantly, the increased DFHBI-1T inside and outside clusters under co-folding condition also revealed that our assay did not inhibit broccoli formation or DFHBI-1T function.

Finally, we examined DFHBI-1T levels in clusters formed by phase-separating separately folded *shu-top* and *shu-bottom* RNAs. While the raw DFHBI-1T level inside of clusters (0.015±0.001) was 8-fold higher than that recorded outside of the clusters (0.002±0.0002), it was even slightly lower to the ones detected for *shu-top* (0.030±0.002) and *shu-bottom* (0.024±0.001), our negative controls (Figure 1e-g). Additionally, after normalization to the RNA intensity (Figure S1a), we observed no significant difference between the DFHBI-1T levels inside and outside the clusters (Figure 1h), which also remained similar to the ones recorded for the negative controls. Thus, these results revealed that no additional broccoli structure formed between *shu-top* and *shu-bottom* when the two separately-folded RNAs phase separated. Collectively, our data revealed that RNA clustering does not require intermolecular base pairing driven by extensive sequence complementarity nor widespread unfolding of the secondary RNA structure.

### 3′ untranslated regions of germ granule mRNAs remain folded upon localization to germ granules

Given our result with *shu-top* and *shu-bottom* RNAs, we reasoned that the absence of stable interactions among mRNAs in RNA clusters in germ granules we reported previously (*36*) might also be due to a well-folded RNA structure. We further hypothesized that, while a specific RNA structure might change during localization, the RNA could still remain well-folded (would retain its structuredness) and exhibit minimal unwinding. To test this hypothesis, we examined the base accessibility to interactions of germ granule mRNAs using dimethyl sulfate mutational profiling with sequencing (DMS-MaPseq) and probed the folded state of an RNA inside and outside of germ granules. DMS preferentially modifies accessible and unpaired adenosines and cytosines, which creates a mutational profile during reverse transcription (*49, 50*).

Exposure to DMS fragments the RNA, making it susceptible to loss during subsequent germ granule purification. In addition, the vitelline membrane surrounding the embryos prevents delivery of DMS. To circumvent these issues, we probed RNA with DMS using isolated germ granules. We purified germ granules marked by a fluorescently-tagged core germ granule protein Vasa (Vasa:GFP) from embryo lysates, as we have done previously (*51, 52*). This approach yielded a pellet with germ granule-bound mRNAs as well as a soluble fraction that contained mRNAs outside the germ granules (Figure S1ci-ii). Importantly, germ granules resisted treatment with 10% DMS (Figure S1cii). Moreover, germ granule mRNAs *nanos* (*nos*), *polar granule component* (*pgc*), and *germ-cell-less* (*gcl*) exhibited 110, 118, and 91-fold enrichment in the pellet relative to the soluble fraction, respectively (Figure S1di-iii), indicating that we were probing the structure of germ granule-bound mRNAs rather than the structure of mRNAs outside the granules.

We specifically focused our studies on the 3′ untranslated regions (UTRs) of *nos*, *pgc* and *gcl* because these regions drive the localization of *nos*, *pgc* and *gcl* to germ granules (*46*). Furthermore, since germ granule mRNAs translate within germ granules (*46, 53–56*) we anticipated that the RNA folding and the intermolecular interactions would be minimally perturbed by ribosomes within the 3′UTRs.

Focusing first on the 3′UTR of unlocalized *nos*, we recapitulated the previously characterized structure of its translational control element (TCE) bound by Smaug protein (Figure 2a) (*57, 58*). This result, together with the fact that DMS efficiently recapitulates the structure of the yeast 18S rRNA to 94% accuracy *in vivo* (*50*), demonstrated the effectiveness of the DMS-MaPseq in probing the base accessibility of germ granule mRNAs in a proteinaceous environment.

**Figure 2:**
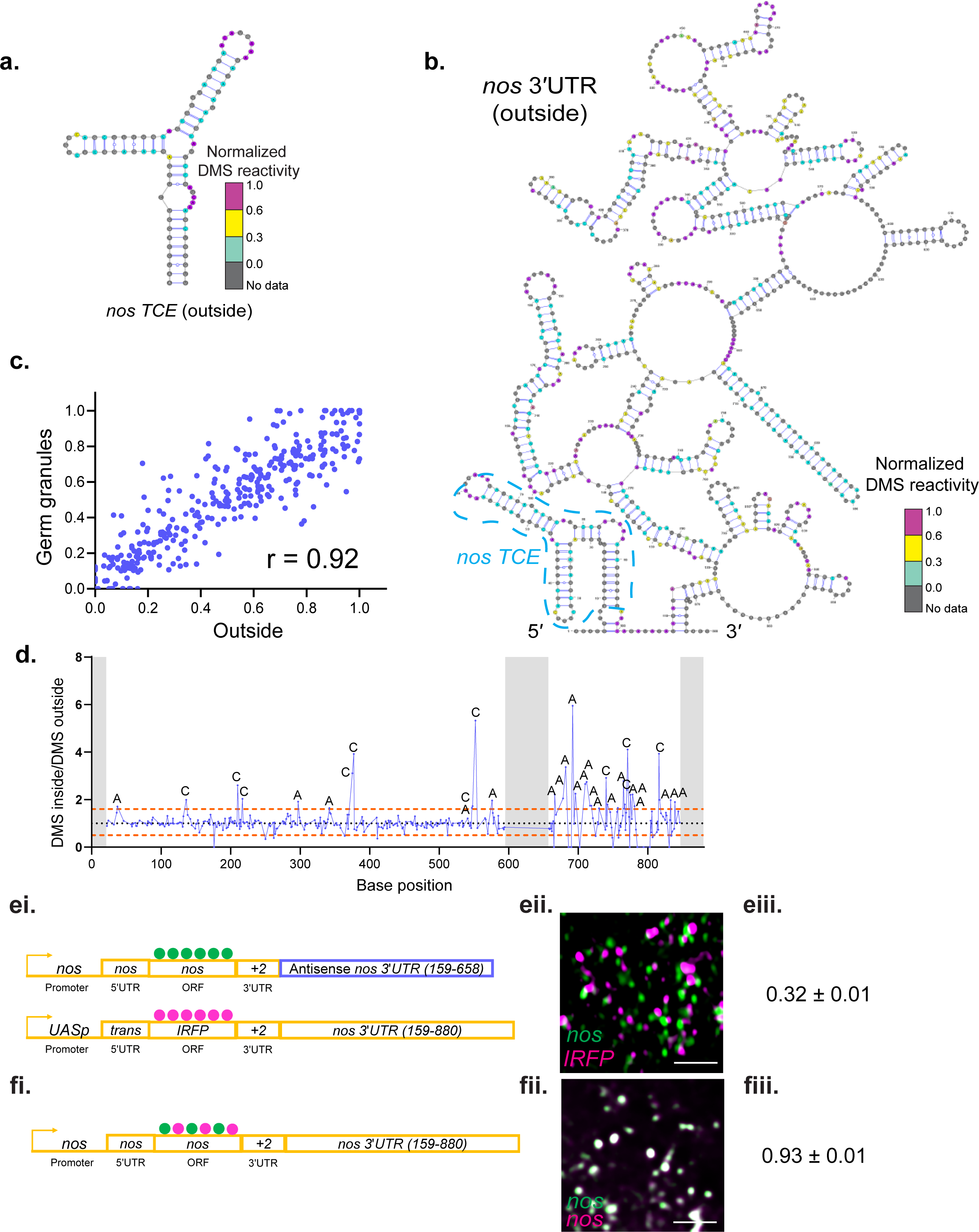
Germ granule mRNAs remain well-folded upon localization to germ granules. **a.** Predicted secondary structure of *nos TCE* outside the germ granules with a mapped normalized DMS reactivity generated by DMS-MaPseq. The normalized DMS reactivities were based on an average of two biological replicates (see Figure S1e-f for the correlation coefficients between the two replicates). **b.** An example of the predicted secondary structure of *nos* 3′UTR outside the germ granules with mapped reactivity by DMS-MaPseq (an average of two replicates is shown). Blue dashed box: *nos TCE*. **c**. Correlation of the normalized DMS reactivities between inside (y-axis) and outside of germ granules (x-axis). Each purple dot represents the DMS reactivity of the same probed bases on *nos* 3′UTR recorded from inside (y-axis) and outside of germ granules (x-axis). **d.** The ratio of the normalized DMS reactivity between inside and outside of the germ granules for the same probed bases on *nos* 3′UTR. Black dotted line: no change in DMS reactivity. Blue dots and line: ratio of DMS reactivities between inside and outside of germ granules for the same probed bases. Orange dashed lines represent thresholds with a p < 0.05, at which the nucleotides are significantly more exposed inside (above 1) or outside the granules (below 1). Grey blocks: no data due to gaps in PCR amplification or primer attachment. The bases, which were significantly more exposed inside the granules, were labeled. **e.** Schematic of an *nos* mRNA fused with a *nos* localization element +2 (*59*) and an antisense *nos* 3′UTR generated by CRISPR/Cas9-PhiC31 approach and of a transgenically-expressed chimeric mRNA containing a CDS of IRFP fused with WT *nos* 3′UTR. The CDS of both mRNAs were hybridized with spectrally distinct smFISH probes (green for *nos* ORF and magenta for *IRFP* ORF) (i). The two mRNAs display low co-localization in germ granules (ii), with a PCC(Costes) of 0.32±0.01. n=15 embryos. Data: Mean±SEM. **f.** Endogenous *nos* hybridized with spectrally-distinct smFISH probes (green and magenta), which alternately hybridized with the *nos* ORF(i), displaying high co-localization (ii), with a PCC(Costes) of 0.93±0.01. n=7 embryos. Data: Mean±SEM. Scale bars: in all images 2 μm. See also Figures S1 and S2, Tables S2 and S3.

We performed two replicates of DMS-MaPseq for RNA fractions from both outside and inside the germ granules. The Pearson correlation coefficients (r) between the two replicates of *nos, pgc* and *gcl* 3′UTRs outside of germ granules were 0.93, 0.97 and 0.99, respectively (Figure S1e), while inside of germ granules they were 0.99, 0.98 and 0.89 for *nos, pgc* and *gcl* 3′UTRs, respectively (Figure S1f). These results demonstrated a high biological reproducibility among DMS-MaPseq experiments. We observed that the 3′UTRs of *nos*, *pgc* and *gcl* outside of germ granules were highly folded and exhibited minimal single-strandedness (Figure 2b, S2a,b). Importantly, the average DMS reactivity of each probed base of *nos*, *pgc,* and *gcl* 3′UTRs inside the germ granules showed remarkable similarity to the average reactivity of the same base recorded for these mRNAs outside the germ granules (correlation coefficients of 0.92, 0.98, 0.82, respectively; Figure 2c, S2a-d, Table S2). Thus, upon localization, *nos*, *pgc,* and *gcl* 3′UTRs remained folded to a similar degree as outside of germ granules (Figure 2c, S2a-d).

Next, we calculated the ratio of DMS signals of mRNAs inside and outside the germ granules to compare the base accessibility of same probed bases. We observed no significant change in DMS reactivity in the *nos* 3′UTR, except for a few nucleotide bases scattered across the 3′UTR and at the AU-rich 3′ end (Figure 2d). We observed a similar result for the *pgc* 3′UTR (Figure S2e). We recorded a more significant change in DMS reactivity in the first 300 nts of the *gcl* 3′UTR (Figure S2fi). However, these changes primarily occurred at the level of individual bases rather than continuous regions (Figure S2fii). These subtle changes in DMS reactivity revealed that the 3′UTR structures of *nos*, *pgc* and *gcl* changed upon localization, suggesting remodeling of the RNA (Figure S3). Notably, despite these changes, the three 3′UTRs remained well-folded within germ granules.

To substantiate our findings further, we co-expressed two distinct mRNAs that exhibited 499 nts of perfect complementary in their 3′UTRs. Both mRNAs contained the +2-localization element derived from *nos* 3′UTR, ensuring localization of both transcripts (*59*) (Figure 2ei). We expected that if these two mRNAs underwent significant structural unfolding in germ granules, then they would base pair and assemble into the same cluster. However, upon co-expression, the two mRNAs enriched in the same granule but formed distinct clusters (PCC(Costes) r: 0.32±0.01, (Figure 2eii-iii); see (*17*) for the description of PCC(Costes)). For comparison, we observed a high co-localization for a doubly-stained WT *nos* mRNA (PCC (Costes) r: 0.93±0.01, (Figure 2fi-iii, Table S3)). Collectively and consistent with our *in vitro shu-top* and *shu-bottom* results, these data demonstrated that *in vivo*, the 3′UTR of mRNAs remained well-folded in germ granules, which limited their ability to engage in base pairing driven by extensive sequence complementarity.

### Base pairing among 3′UTRs of germ granule mRNAs is driven by regions with low sequence complementarity and exhibits low probability for sustained interactions

Our experiments revealed that mRNAs remain well-folded in the RNA clusters of germ granules (Figure 1, 2). However, the ability of RNAs to condense in the absence of other cellular components *in vitro* (Figure 1c; (*16, 28, 29, 60–62*)) nevertheless highlights their capacity for intermolecular interactions that are challenging to directly examine through microcopy, *in vitro* phase separation assays and DMA-MaPseq probing. To further understand the abundance and the properties of intermolecular base pairing in RNA clusters further, we simulated tertiary structures of *nos, pgc,* and *gcl* 3′UTRs and their homodimers using our previously established minimal coarse-grained SIS model (*34*). Importantly, our simulation reproduced the known structure of the *HIV* U4/6 core region (Figure S4a-c; (*63*)), which established the SIS model as a tool to investigate three-dimensional (3D) RNA folding.

We first modeled the 3D structures of *nos, pgc* and *gcl* 3′UTR monomers. We observed that *nos, pgc* and *gcl* 3′UTRs adopt many distinct and highly folded secondary structures with different energy states (Figure 3a,b, S4d-g) revealing the structural heterogeneity of these RNAs. Moreover, we determined that regardless of the energy states, most secondary structures of *nos, pgc* and *gcl* 3′UTR monomers displayed long and stable stems with numerous loops engaged in intramolecular base pairing driving tertiary folding (Figure 3a,b, S4d-g). This finding is consistent with the ensemble measurements of base accessibility obtained by DMS-MaPseq which revealed that unlocalized *nos, pgc* and *gcl* 3′UTRs are well-folded (Figure 2b, S2a-b).

**Figure 3:**
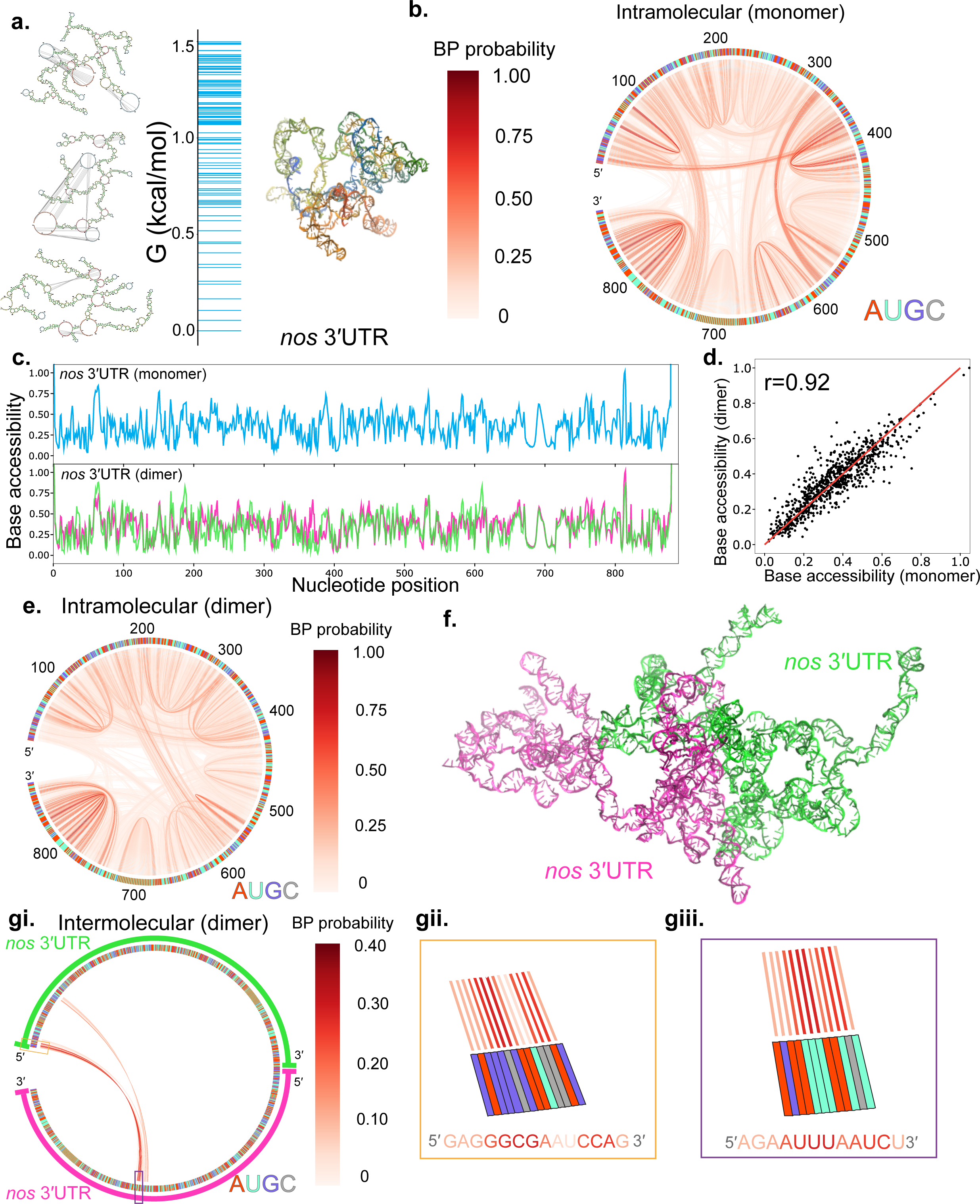
Base pairing within *nos* 3′UTR homodimer is driven by regions with low sequence complementarity and exhibits low probability for sustained interactions. **a.** Examples of simulated structures of monomeric *nos* 3′UTR at different energy states with an accompanying free energy spectrum where each blue line represents a cluster of secondary structures generated in the simulation. Shown is a 3D conformation of *nos* 3′UTR corresponding to the ground state (G = 0 kcal/mol). **b**. Intramolecular base pairing (red lines) within *nos* 3′UTR in a monomer state. **c**. Simulated base accessibility of each nucleotide of *nos* 3′UTR in a monomer state (top; blue line) and homodimer state (bottom; magenta and green lines). **d.** Correlation of base accessibility between homodimer and monomer state of *nos* 3′UTR, with a Pearson correlation coefficient (r) of 0.92. **e.** Intramolecular base pairing (red lines) within *nos* 3′UTR in a homodimer state. **f.** Simulated *nos* 3′UTR homodimer (magenta and green). **g.** Intermolecular base pairing (red lines) between two *nos* 3′UTRs (i) with zoomed-in regions of interacting regions (ii, iii). See also Figures S3-S5.

Next, we simulated intermolecular base pairing for *nos, pgc* and *gcl* 3′UTR homodimers. Here, a homodimer refers to two RNA molecules that interacted with each other using one or more base pairs without a predefined structure, interaction strength, duration, or the involvement of a particular RNA sequence. We assumed RNA concentrations for *nos, pgc* and *gcl* 3′UTRs of 10.7µM, 29.4 and 50.7 µM, respectively, which were similar to the concentrations reported for these RNAs within the RNA clusters *in vivo* (*36*). Notably, we detected minimal changes in base accessibility between monomeric and homodimeric states of *nos, pgc* and *gcl* 3′UTRs (r = 0.92, 0.73 and 0.81, respectively) (Figure 3c,d, S4h-k), which implied the absence of extensive structural melting upon homodimerization. These data demonstrated that within homodimers RNAs remained well-folded while retaining their ability to interact with the neighboring RNAs. In support of this observation, 99%, 95% and 97% of the base pairing within homodimers remained intramolecular for *nos, pgc* and *gcl* 3′UTR, respectively, revealing controlled intermolecular base pairing upon dimerization (Figure 3e,f, S5a-d).

We further observed that intermolecular base pairing within the *nos, pgc* and *gcl* 3′UTR homodimers was predominantly driven by scattered and discontinuous stretches of bases, resulting from regions of low sequence complementarity (Figure 3gi-iii, Figure S5ei-fiii). Specifically, in *nos* 3′UTR homodimers, 5 out of 22 interacting bases at the 5′ end of *nos* 3′UTR had a base pairing probability close to 0.4, while the remaining interacting bases had a base pairing probability of ∼ 0.1 (Figure 3gi). Similar results were also observed in *pgc* and *gcl* 3′UTR homodimers (Figure S5ei,fi).

In contrast to *osk, bcd* and *HIV* which use a single GC-rich complementary sequence (CSs) to establish stable dimerization (66% (*bcd*) (*39, 44*) to 100 % (*osk*, *HIV*)) recoverable by the RNA gel (*38, 40–43*), none of the sequences that engaged in intermolecular base pairing of *nos, pgc* and *gcl* 3′UTRs resembled the GC-rich CSs found on *osk, bcd* and *HIV* (Figure 3gi-giii, S5ei-fiii). To determine whether the CSs on *osk, bcd* and *HIV* are specially designed to support a sustained RNA dimerization or if any CSs of similar length would achieve the same outcome, we modified the *HIV* SL1 stem-loop and replaced its dimerization CS with CSs of the same length but varying GC content. We then inserted these modified stem-loops into the *shu* 3′ UTR, which is a dimerization inert RNA (Figure 1b, S6ai,ii). After *in vitro* transcription, we evaluated the dimerization of RNAs on a non-denaturing agarose gel following established protocols (*39, 41*). We observed strong dimerization only for CSs with 100% GC content (Figure S6ai,ii, Table S1). Among the three tested CSs with 66% GC content, only one (GUGCAC) dimerized, while others, including those with lower GC contents, did not (Figure S6ai,ii, Table S1). In addition, the concatenation of shorter (< 6nts) CSs with 100% GC content or AU-rich CSs, with a length of 14 nts, which is longer than any of the interacting AU-rich regions on *nos, pgc* and *gcl* 3′UTRs (Figure 3gi, S6ei, fi), did not promote stable dimerization (Figure S6b,c, Table S1). These data indicated that the length and GC content of a CS were the primary determinants of sustained intermolecular base pairing *in vitro*. Consistent with these observations, from our simulations, none of the *nos, pgc* and *gcl* 3′UTRs sequences engaged in intermolecular base pairing were driven by GC-rich CSs found in *osk, bcd* and *HIV* or their dimerization-competent variants (Figure 3g, S5e,f, S6a).

Together, our *in silico* experiments revealed that intermolecular base pairing within *nos, pgc* and *gcl* 3′UTR homodimers is driven by regions with low complementarity rather than by specific RNA sequences and exhibits a low probability for sustained interactions. Additionally, while engaging in intermolecular base pairing, the RNA remained folded, supporting our *in vitro* and *in vivo* findings (Figure 1, 2, 3g, S5e,f) (*36*).

### Exposed GC-rich complementary sequences enhance interactions among engineered *nos* mRNAs *in vivo*

Intermolecular base pairing observed for clustered germ granule mRNAs was independent of specific RNA sequences, distinguishing it from the stable dimerization observed in *osk, bcd* and *HIV,* which results from exposed GC-rich CSs (*36, 38–44*). To examine the effect of exposed GC-rich CSs on germ granule mRNAs, we created RNA constructs containing four stems derived from the *HIV* SL1 dimerization sequence (*43*) where each stem contained GC-rich CSs derived from the *HIV* or *nos* 3′UTR (see Figure 7c, green lines) (termed *HIV* and *nos*, respectively). As a control, we also generated a construct that contained a non-complementary sequence in the loop (termed *non*) (Figure S7ai). Using *in vitro* transcribed RNAs and non-denaturing agarose gels, we observed an enhanced dimerization of *HIV* and *nos* but not of *non* RNA (Figure S7aii, Table S1). These results were consistent with the prediction that only GC-rich CSs dimerize RNAs (Figure S6ai, ii).

To test whether *HIV* and *nos* CSs enhance intermolecular interactions of mRNAs *in vivo*, we inserted these constructs after the non-coding *LaczA* and *LaczB* RNA sequences (*64*) and examined intermolecular interactions using single-molecule fluorescent in situ hybridization (smFISH) and co-localization analysis in *Drosophila* Ras cells (Figure S7b-d). To verify that the intermolecular interactions were not unique to *HIV* or *nos* CSs, we also inserted GC-rich *sense* and *antisense* sequences within the loops of *LaczA* and *LaczB* RNAs (Figure S7e).

Applying Pearson’s Correlation Coefficient (PCC) approach, which reflects the linear relationship between intensities of fluorescently-labeled *LaczA* and *LaczB* mRNAs, we demonstrated that *LaczA* and *LaczB* that contained *HIV, nos* and *sense-antisense* CSs better co-localized (r(PCC): 0.57±0.03, 0.50±0.02, and 0.49±0.03, respectively) than those that contained *non* sequences (r(PCC):0.33±0.02) (Figure S7f). Importantly, this co-localization could not be explained by differential expressions among the constructs (Figure S7g-h). Instead, these data confirmed that exposed GC-rich CSs promoted interactions among *LaczA* and *LaczB* mRNAs in cells.

To then examine the effect of exposed GC-rich CSs in flies, we inserted the *HIV*, *nos* and *non* constructs into 3′UTR of *nos* gene using CRISPR/Cas9-PhiC31 approach (termed *nos-HIV, nos-nos* and *nos-non,* respectively) (Figure 4a). After eight rounds of crosses with balancer flies to remove possible off-target effects induced by guide RNAs, we crossed *nos-non, nos-HIV,* and *nos-nos* flies with *nos^BN^* flies, which limited expression of WT *nos* to early oogenesis (*65*), and thus allowed examination of only the edited *nos* alleles during embryogenesis. Using DNA gel electrophoresis and the cDNA products extracted from eggs laid by *nos-non/nos^BN^, nos-HIV/nos^BN^,* and *nos-nos/nos^BN^* females, we determined that the modified *nos* transcripts were spliced normally (Figure 4bi-bii, Table S4). In addition, qRT-PCR analysis revealed that the 3′UTR processing of the *nos-non, nos-HIV,* and *nos-nos* mRNAs was similar (Figure 4ci-cii, Table S4).

**Figure 4:**
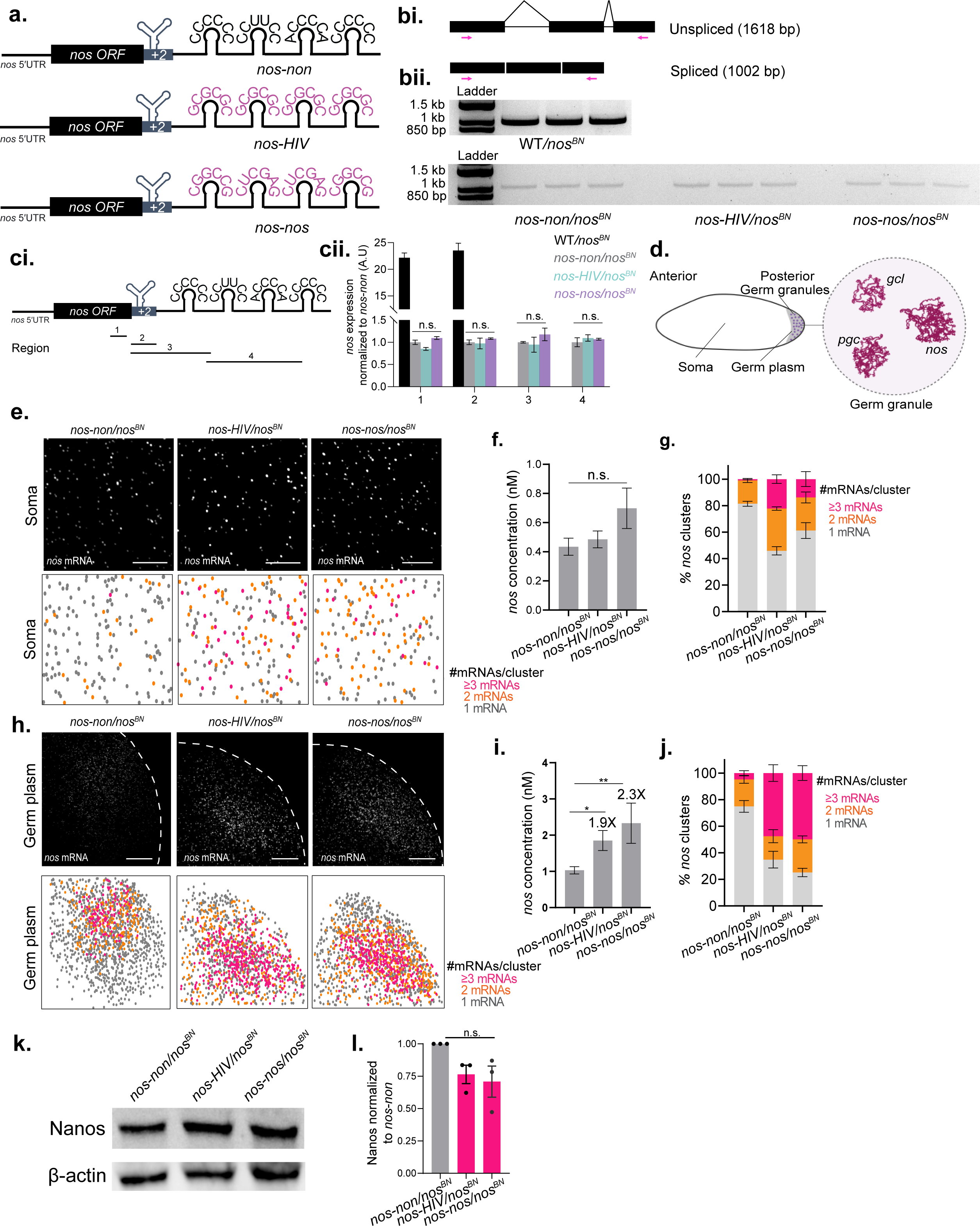
Exposed GC-rich complementary sequences enhance interactions among engineered *nos* mRNAs *in vivo.* **a.** Schematics of the endogenous *nos* containing four stems with non-palindromic (*nos-non*), *HIV-* (*nos-HIV*) or *nos-*derived (*nos-nos*) palindromes in its 3′UTR. **b.** Schematics of unspliced or spliced *nos* transcript where the primer targeted regions were shown in magenta arrows (i). *nos* PCR products from the cDNAs (100 ng total input) of the embryos laid by WT*/nos^BN^* (top)*, nos-non/nos^BN^*, *nos-HIV/nos^BN^* and *nos-nos/nos^BN^*flies (bottom). Each genotype had three lanes shown which represented three replicates. **c.** Schematic of the primer targeted regions (1, 2, 3 and 4) on *nos*-*non*, which are also used for WT *nos*, *nos-HIV* and *nos-nos* (i). The expression levels of each region are determined by qRT-PCR (ii). n = 3 for all genotypes. Data: Mean±SEM. n.s.: not statistically significant (unpaired t-test). **d.** Schematic of the *Drosophila* embryo and mRNA clusters in its germ granules. **e.** smFISH of *nos-non, nos-HIV,* and *nos-nos* mRNAs in soma of the embryos laid by *nos-non/nos^BN^*, *nos-HIV/nos^BN^* and *nos-nos/nos^BN^* females (top). Heat maps were generated based on the number of mRNAs per cluster (bottom). Scale bars: 4 μm. **f.** Concentrations of *nos-non* (n=7), *nos-HIV* (n=6) and *nos-nos* (n=6) mRNAs in soma. Data: Mean±SEM. n.s.: not statistically significant (unpaired t-test). **g.** Number of *nos* mRNAs per cluster in soma of the embryos laid by *nos-non/nos^BN^* (n=7), *nos-HIV/nos^BN^* (n=6) and *nos-nos/nos^BN^* (n=6) flies. Data: Mean±SEM. **h.** smFISH of *nos-non, nos-HIV,* and *nos-nos* mRNAs in the germ plasm of the embryos laid by *nos-non/nos^BN^*, *nos-HIV/nos^BN^*and *nos-nos/nos^BN^* females (top). Heat maps were generated based on the number of mRNAs per cluster (bottom). Scale bars: 17 μm. **i.** Concentrations of *nos-non* (n=6), *nos-HIV* (n=6) and *nos-nos* (n=6) mRNAs in the germ plasm. Data: Mean±SEM. n.s.: not statistically significant (unpaired t-test). **j.** Number of *nos* mRNAs per cluster in the germ plasm of the embryos laid by *nos-non/nos^BN^* (n=7), *nos-HIV/nos^BN^* (n=6) and *nos-nos/nos^BN^* (n=6) flies. Data: Mean±SEM. **k.** Example western blot of anti-Nanos and anti-β-actin (loading control) from the embryos laid by *nos-non/nos^BN^*, *nos-HIV/nos^BN^* and *nos-nos/nos^BN^* flies. **l.** Quantification of Nanos protein levels normalized to β-actin and *nos-non*. Three biological replicates were used. Data: Mean±SEM. n.s: not significant. See Figure S6 and Table S4.

However, compared to WT *nos* expressed in WT/*nos^BN^*eggs, the mRNA levels of *nos-non*, *nos-HIV*, and *nos-nos* were reduced by approximately 22-fold (Figure 4bii; 4ci-cii). These data revealed that the changes to the 3′UTR interfered with the expression of *nos* mRNA (see Figure 4a and Methods). This reduction was anticipated given that GC-rich sequences and strong RNA structures located into the 3′UTR trigger RNA degradation (*66, 67*). Therefore, in our experiments, the phenotypes of *nos-non* served as the baseline against which we evaluated the effects of *HIV* and *nos* GC-rich CSs on the gene expression of the engineered *nos* mRNAs.

Using smFISH, we first examined the clustering ability of *nos-non, nos-HIV,* and *nos-nos* mRNAs in embryos. We determined previously that WT *nos* forms clusters only in germ granules (Figure 4d) (*17, 18, 36, 37*). However, only 45± 3.0% *nos-HIV* and 60±6.0% *nos-nos* mRNAs appeared as single mRNAs outside of germ granules compared to 81±1.9% detected for *nos*-*non* mRNA (Figure 4e-g). Notably, the somatic expression of the three *nos* constructs was comparable (Figure 4f), indicating that the somatic increase in mRNA clustering of *nos-HIV* and *nos-nos* was not due to differences in mRNA concentration.

In addition, *nos-HIV* and *nos-nos* mRNAs exhibited a 1.9 and 2.3-fold higher enrichment in the germ plasm, respectively, compared to *nos-non* mRNAs (Figure 4h-j). Moreover, only 35±6.3% and 25±3.2% for *nos-HIV* and *nos-nos* mRNAs respectively were single transcripts in germ plasm compared to 75±4.4% detected for *nos-non* mRNAs (Figure 4j). Additionally, western blot analysis revealed that *nos-non/nos^BN^*, *nos-HIV/nos^BN^* and *nos-nos/nos^BN^* exhibited similar Nanos protein expression (Figure 4k, l).

Together these data demonstrated that compared to the WT *nos*, *nos-non, nos-HIV* and *nos-nos* experienced a significant reduction in their gene expression. However, compared with each other, *nos-non*, *nos-HIV* and *nos-nos* mRNA had similar splicing and 3′UTR processing efficiencies, efficiently localized to germ granules and produced similar levels of Nanos protein.

### HIV-derived SL1 stem loop in *nos* 3′UTR impairs fly embryogenesis

Nanos protein is essential for abdominal patterning and egg hatching of the embryo (*68, 69*). To examined the effect of the 22-fold reduction in gene expression recorded for the *nos* CRISPR alleles on fly embryogenesis (Figure 4cii), we counted the body segments using smFISH on *fushi tarazu* mRNAs and examined egg hatching rates as described previously (*51, 70*). We observed that control eggs laid by WT/*nos^BN^* females formed seven (WT) abdominal segments and displayed egg hatching rates of 86.2%±1.6% (Figure 5a-c), consistent with the phenotype recorded for WT flies (*51*). In contrast, eggs laid by *nos*-*non/nos^BN^*females formed on average 3.8±0.1 abdominal segments (Figure 5b) and displayed the egg hatching rate of 3%±0.7% (n=2280 eggs) (Figure 5c). Therefore, the insertion of HIV-derived SL1 stem loops into the *nos* 3′UTR not only reduced expression of *nos* mRNA (Figure 4cii) but also severely impaired fly embryogenesis.

**Figure 5:**
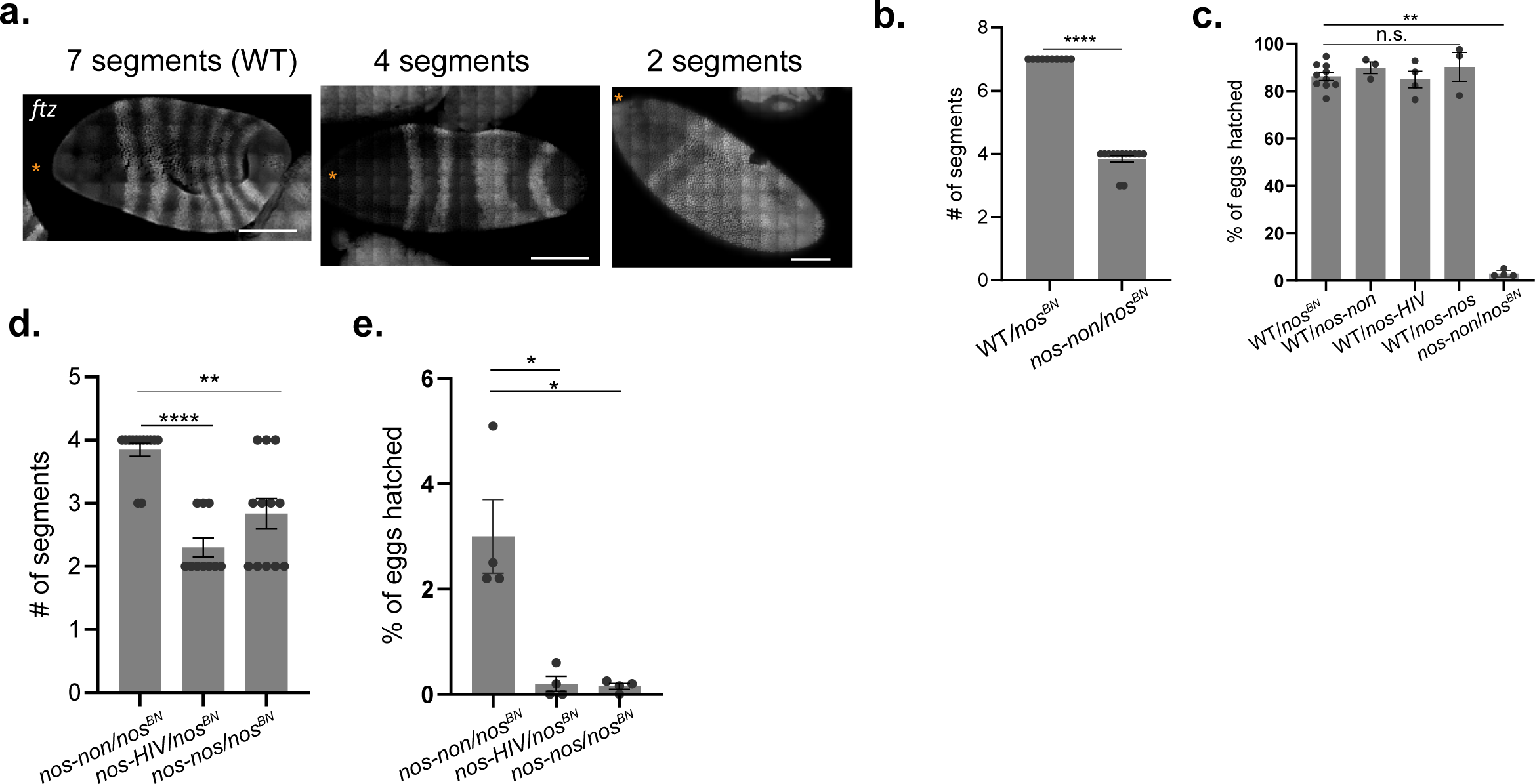
HIV-derived SL1 stem loop and exposed GC-rich complementary sequences impair fly embryogenesis. **a.** Body segments stained by *ftz* mRNA in embryos. Asterisks: anterior tips of the embryos. Scale bars: 100 μm. **b.** Average number of body segments in the embryos laid by WT*/nos^BN^* (n=10) and *nos-non/nos^BN^* (n=13). Data: Mean±SEM. ****: p < 0.0001 (unpaired t-test). **c.** Percentages of eggs hatched in WT/*nos^BN^* (n=1951 eggs), WT/*nos-non* (n=880), WT/*nos-HIV* (n=2136 eggs), WT/*nos-nos* (n=643 eggs) and *nos-non*/*nos^BN^* (n=2280 eggs). At least three separate counting experiments were performed for each condition. Data: Mean±SEM. n.s: not significant. **: p < 0.01 (unpaired t-test). **d.** Average number of body segments in the embryos laid by *nos-non/nos^BN^* (n=13)*, nos-HIV/nos^BN^* (n=10) and *nos-nos/nos^BN^* (n=12) females. Note that *nos-non/nos^BN^* data was the same as in **b.** **, ****: p < 0.01 and 0.0001, respectively (unpaired t-test). **e.** Percentages of eggs hatched in *nos-non*/*nos^BN^* (n=2280), *nos-HIV*/*nos^BN^* (n=2568) and *nos-nos*/*nos^BN^* (n=1698). Note that *nos-non/nos^BN^* data was the same as in **c.** At least three separate counting experiments were performed for each condition. Data: Mean±SEM. n.s: not significant. *: p < 0.05 (unpaired t-test).

### Exposed GC-rich complementary sequences exacerbate the embryogenesis phenotypes induces by the HIV-derived SL1 stem loop

*nos-non*, *nos-HIV* and *nos-nos* mRNAs had similar splicing and 3′UTR processing efficiencies, expressed at similar levels, and accumulated a similar amount of Nanos protein (Figure 4). However, we also observed that eggs laid by *nos-HIV/nos^BN^* and *nos-nos/nos^BN^* females formed 2.3±0.2, and 2.8±0.2 segments per embryo, respectively, which was significantly lower than *nos*-*non/nos^BN^* embryos (Figure 5d). In addition, the *nos-HIV/nos^BN^* and *nos-nos/nos^BN^*embryos displayed egg hatching rates of 0.20±0.14% (n=2568 eggs) and 0.15±0.05% (n=1698 eggs), respectively, which were 15- and 20-times lower compared to *nos*-*non/nos^BN^* embryos (Figure 5e).

These exacerbated segmentation phenotypes cannot be explained by differences in Nanos protein levels, as all CRISPR lines produced similar amount of protein (Figure 4k, l) nor by de-repression of unlocalized *nos,* which would support normal abdominal segmentation (*71*). Instead, these results indicate that an alternative mechanism, specifically sensing the exposed GC-rich sequences triggers these phenotypes. Together our data demonstrated that while the HIV-derived SL1 stem loops produced the dominant embryogenesis defect, the exposed GC-rich CSs exacerbated it independent of gene expression mechanism.

### HIV-derived stem loop and exposed GC-rich complementary sequences impair fly oogenesis

Nanos protein is also required for the maintenance of germline stem cells (GSCs) during oogenesis (*72, 73*). To investigate whether *nos-non*, *nos-HIV* and *nos-nos* functioned similarly in oogenesis, we crossed *nos-HIV*, *nos-nos,* and *nos-non* flies with *nos*-deficient flies (*nos^def^*), which lacked the *nos* gene, allowing us to investigate the expression of edited *nos* alleles during oogenesis in *nos-HIV/nos^def^*, *nos-nos/nos^def^,* and *nos-non/nos^def^*females. We compared the ovary morphology of these females to their sibling controls (termed *nos-HIV/*WT*, nos-nos/*WT*, nos-non/*WT, respectively) at 3, 9, and 14 days after females were enclosed. While siblings formed round and fecund ovaries (Figure 6a, S8ai-aiii), ovaries of *nos-non*/*nos^def^* females exhibited an intermediate phenotype, with 69%, 72% and 71% of the examined ovaries being smaller than the ovaries of siblings aged 3, 9 and 14 days (Figure S8b). As noted for embryos, these defects in oogenesis were anticipated given the reduced expression of CRISPR alleles compared to WT *nos* (Figure 4cii).

**Figure 6.**
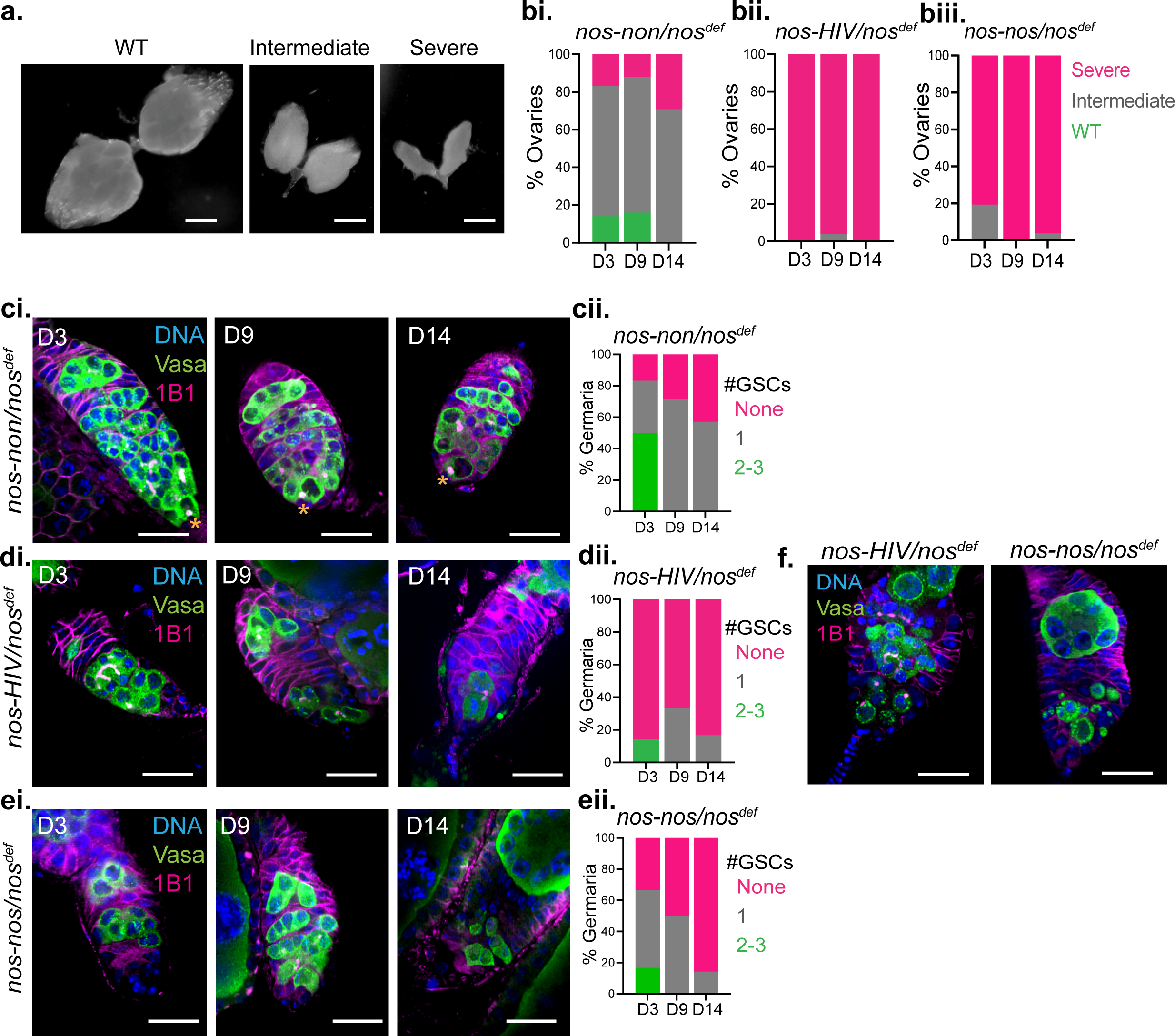
HIV-derived SL1 stem loop and exposed GC-rich complementary sequences impair fly embryogenesis. **a.** Images of WT, intermediate and severe phenotypes of ovary morphology. Scale bars: 200 μm **b.** Percentages of examined ovaries with a WT, intermediate and severe phenotypes in *nos-non*/*nos^def^* (n=29, 25 and 24) (i), *nos-HIV*/*nos^def^* (n=29, 26 and 22) (ii) and *nos-nos*/*nos^def^* (n=31, 23 and 26) (iii) females aged 3, 9 and 14 days (D3, D9 and D14, respectively). **c.** Immunofluorescent images of germaria of *nos*-*non*/*nos^def^* (i). Asterisks: GSCs. Blue: DNA/DAPI; green: germ cells immunostained with anti-Vasa antibody; magenta: spectrosomes immunostained with anti-1B1 antibody. Scale bars: 18 μm. Percent of germaria with 0, 1 or 2-3 GSCs in *nos-non*/*nos^def^* (ii). n = 6, 7 and 7 germaria for D3, D9 and D14, respectively. **d.** Immunofluorescent images of germaria of *nos*-*HIV*/*nos^def^*(i). Asterisks: GSCs. Blue: DNA/DAPI; green: germ cells immunostained with anti-Vasa antibody; magenta: spectrosomes immunostained with anti-1B1 antibody. Scale bars: 18 μm. Percent of germaria with 0, 1 or 2-3 GSCs in *nos-HIV*/*nos^def^* (ii). n = 7, 7 and 7 germaria for D3, D9 and D14, respectively. **e.** Immunofluorescent images of germaria of *nos*-*nos*/*nos^def^* (i). Asterisks: GSCs. Blue: DNA/DAPI; green: germ cells immunostained with anti-Vasa antibody; magenta: spectrosomes immunostained with anti-1B1 antibody. Scale bars: 18 μm. Percent of germaria with 0, 1 or 2-3 GSCs in *nos-nos*/*nos^def^* (ii). n = 6, 6 and 7 germaria for D3, D9 and D14, respectively. **f.** Aberrant germ cell morphology (green) in germaria derived from *nos*-*HIV*/*nos^def^*and *nos*-*nos*/*nos^def^* females. Scale bars: 18 μm.

Strikingly, 100% and 81% of ovaries from *nos-HIV/nos^def^*and *nos-nos/nos^def^* 3-day-old females already displayed severe morphological defects, respectively (Figure 6bii, biii). Apart from their much-reduced size, these ovaries also displayed visible abnormalities in oogenesis progression (Figure 6a, b). Importantly, the progression through oogenesis in *nos-HIV/*WT and *nos-nos/*WT females was normal (Figure S8aii, aiii), indicating that the observed phenotypes were not dominant negative.

Moreover, while 83%, 71% and 57% of *nos-non/nos^def^*germaria had at least 1 germline stem cell (GSC) (Figure 6ci, ii, S8bi, ii; cells marked with asterisks), 86%, 67%, and 83% *nos*-*HIV*/*nos^def^* germaria showed a significant depletion of GSCs in 3, 9- and 14-day old females, respectively (Figure 6di, ii, S8ci, cii; cells marked with asterisks). Similarly, *nos-nos/nos^def^* germaria displayed an attenuated GSC depletion compared to the *nos-non/nos^def^* germaria and the *nos-nos/*WT sibling controls (Figure 6ei, ii, S8di,ii; cells marked with asterisks). In addition, some *nos*-*HIV*/*nos^def^* and *nos-nos/nos^def^* germaria also presented aberrant germ cell morphology (Figure 6f). Notably, these exacerbated phenotypes observed in *nos*-*HIV*/*nos^def^* and *nos-nos/nos^def^* were consistent with those recorded in flies lacking Nanos expression (*72, 73*). Together these data indicated that HIV-derived SL1 stem loops severely stunt progression through oogenesis and that exposed GC-rich CSs exacerbated these phenotypes, the consistent with the observations recorded for the embryo.

### GC-rich complementary sequences are embedded within the RNA structure in germ granule mRNAs

The additional developmental phenotypes triggered by exposed GC-rich CSs prompted us to examine the prevalence of these sequences in germ granule mRNAs. CSs consist of palindromes, inverted repeats (IRs) and sense-antisense sequences (Figure 7a), which can base pair intermolecularly or intramolecularly to form RNA structures (Figure 7a).

**Figure 7.**
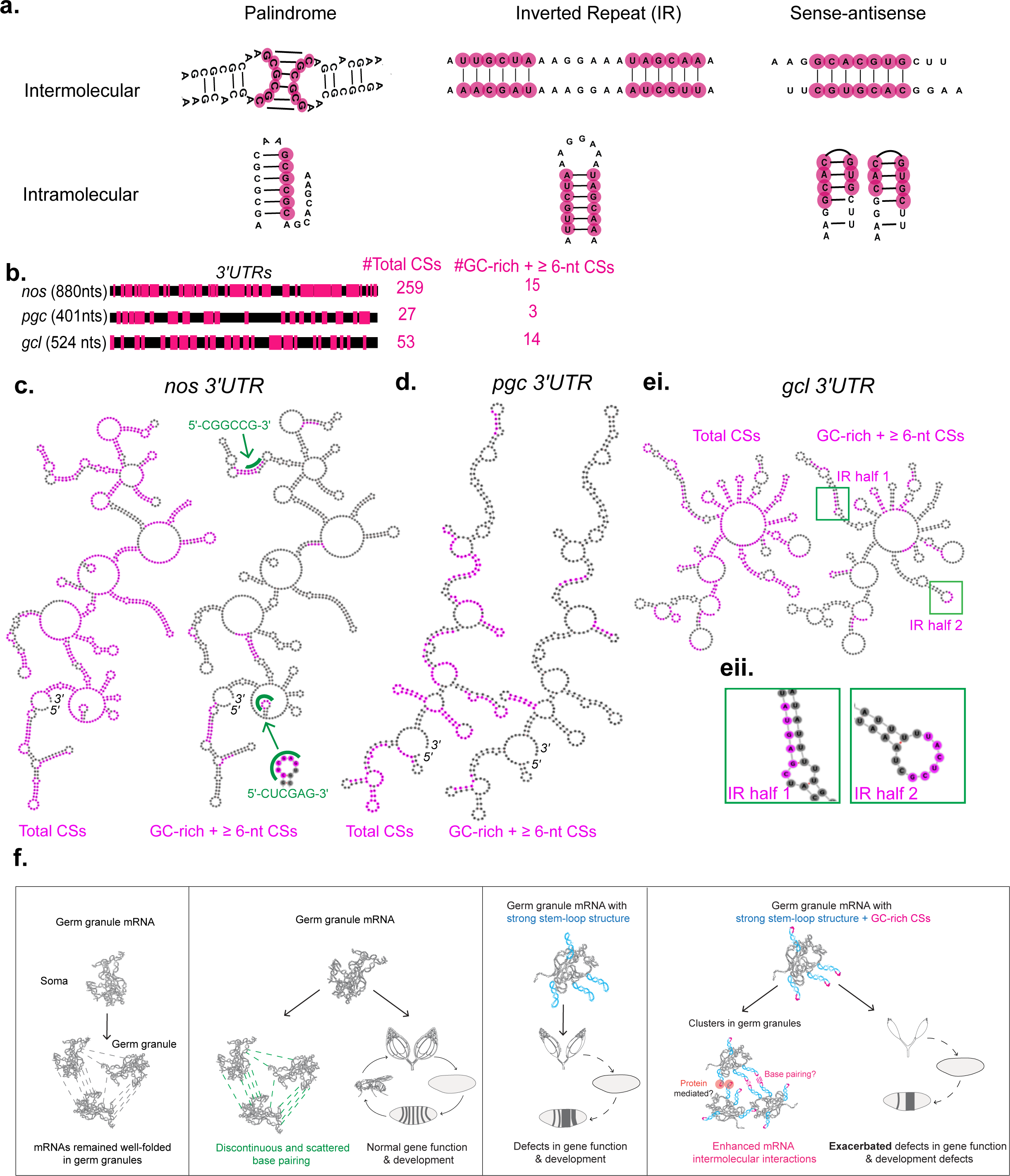
GC-rich complementary sequences are embedded within the RNA structure in germ granule mRNAs. **a.** Types of CSs that enable intermolecular and intramolecular base pairing between mRNAs. **b.** Alignment and abundance of total (magenta boxes) and GC-rich CSs, which had a minimal length of 6 nts in *nos, pgc,* and *gcl* 3′UTRs. **c.** Predicted secondary structures of *nos* 3′UTR with mapped total CSs (magenta, left) or GC-rich CSs with a minimum of 6 nucleotides (magenta, right) outside of the germ granules. The RNA structures were generated based on two DMS-MaPseq replicates. The green sequences (“CGGCCG” and “CUCGAG”) and lines were *nos* CSs used in *nos-nos* experiments. A zoom-in image of “CUCGAG” demonstrating that this sequence is partially embedded within the RNA structure. **d.** Predicted secondary structures of *pgc* 3′UTR with mapped total CSs (magenta, left) or GC-rich CSs with a minimum of 6 nucleotides (magenta, right) outside of the germ granules. The RNA structures were generated based on two DMS-MaPseq replicates. **e.** Predicted secondary structures of *gcl* 3′UTR with mapped total CSs (magenta, left) or GC-rich CSs with a minimum of 6 nucleotides (magenta, right) outside of the germ granules (i). An example of IR in which half of the sequence was base paired intramolecularly (ii). The RNA structures were generated based on two DMS-MaPseq replicates. **f.** Proposed model depicting that mRNAs remain folded within mRNA clusters in germ granules (first panel). The interactions among germ granule mRNAs in clusters are controlled by RNA folding. These interactions are mainly driven by regions with low complementarity and exhibit low probability for sustained interactions (second panel). HIV-derived SL1 stem loops (blue) are the main driver of developmental the defects in fly embryogenesis and oogenesis (panel 3). Notably, exposed GC-rich CSs (red) within SL1 structures exacerbated these phenotypes and also enhanced intermolecular interactions among the mRNAs (panel 4). However, the mechanisms responsible for these developmental defects are unclear. See also Tables S5-S7.

We set out to identify CSs that were similar to those that drive the dimerization of *osk, bcd* and *HIV* RNAs (*38–44*). Since these three RNAs employ a palindrome of 6-nt (*osk*, *HIV*) (*38, 40–43*) or an IR of 12 nt (*bcd*) (*39, 44*) and given that only GC-rich CS drive stable dimerization (Figure S6ai,ii), we limited our search to CSs with a minimal length of 6 nts and 50% GC-content.

Our analysis revealed that the 3′UTRs of *nos*, *pgc*, and *gcl* were replete with CSs. Specifically, they contained 259, 27, and 53 CSs with a minimum length of 6 nts, respectively, of which 15, 3 and 14 had a GC content larger than 50%, respectively (Figure 7b, Tables S5-S7). However, mapping of these CSs onto the 3′UTR structures of unlocalized *nos*, *pgc*, and *gcl* determined by the DMS-MaPseq (Figure 2b, S2a, b) revealed that all GC-rich CSs were embedded within the RNA structure (Figure 7c-ei). Frequently, a part of a CS was exposed in the RNA loop, whereas the rest was base paired intramolecularly (Figure 7eii), rendering these CSs incompetent for intermolecular base pairing. Notably, the CGGCCG of *nos* used in *nos-nos* was completely embedded within the WT *nos* 3′UTR structure while CUGGAG was partially embedded (Figure 7c; green lines).

Together, our analysis revealed that germ granule mRNAs contain many GC-rich CSs competent of stable intermolecular base pairing *in vitro*. However, *in vivo,* these sequences were embedded within the RNA structure which rendered them inaccessible to intermolecular interactions. Thus, our data revealed that RNA structure shields germ granule mRNAs against potential detrimental effects of exposed GC-rich CSs thereby preserving normal mRNA function and fly development.

## Discussion

In this study, we investigated the abundance and type of intermolecular base pairing in *Drosophila* germ granules. Within these granules, mRNAs self-organize into RNA clusters containing multiple transcripts derived from the same gene (Figure 4d) (*17, 18, 36, 37*).

Using *in vitro, in vivo* and *in silico* assays we report that mRNAs remain well-folded upon the formation of RNA clusters (Figure 1, 2,7f). Within the clusters, intermolecular base pairing is driven by scattered and discontinuous stretches of bases, resulting from regions of low sequence complementarity (Figure 3g, S5e, f, 7f). These findings are consistent with the lack of dependence of RNA clustering on a particular RNA sequence we reported previously (*36*). Finally, engineered germ granule mRNAs with exposed GC-rich CSs presented within stable stem loops induce persistent base pairing *in vitro* (Figure S7a) and enhanced intermolecular interactions *in vivo* (Figure 4g, S7b-e) but also disrupt normal fly development (Figure 5, 6, S8, 7f). Notably, while the presence of a strong SL1 stem loop derived from HIV is the major driver of developmental defects (Figure 5, 6, S8, 7f), the exposed GC-rich CSs exacerbate them (Figure 5, 6). Notably, germ granule mRNAs contain numerous GC-rich CSs (Figure 7b), however *in vivo*, these sequences are shielded within the RNA structure (Figure 7c-eii). Thus, our findings underscore the protective role of RNA folding in mitigating the potentially adverse effects of exposed GC-rich CSs and preserving mRNA functionality within environments with increased RNA density, such as RNA granules (Figure 7f).

### Base pairing that exhibits low probability for sustained interactions could enable mobility of germ granule mRNAs

Our *in silico* simulations show that the 3′UTRs of germ granule mRNAs adopt various secondary conformations (Figure 3a,b, S4d-g), thus contributing to the diversity of tertiary folding. This is consistent with recent observations on the *in vitro* transcribed adenosylcobalamin riboswitch aptamer domain, where a single RNA sequence folds into diverse 3D structures (*74*).

An important implication of this finding is that conformational heterogeneity enables multiple interaction combinations among mRNAs (Figure 3g, S5e-f). The multitude of these combinations accommodates the low probability nature of any specific intermolecular base pairing throughout the RNA, while simultaneously maintaining stable RNA structures that prevent sustained intermolecular base pairing. This mode of interaction stands in direct contrast to the one observed in clusters composed of RNAs with repeat sequences, where RNAs within clusters readily unfold and expose single-stranded regions (*75*), accommodating extended intermolecular base pairing (*34*).

Therefore, the balance between the strength and frequency of intermolecular base pairing may be crucial for preventing the formation of pathological RNA aggregates observed in RNAs with expanded nucleotide repeats (*16, 34*) while enabling mRNAs to congregate and maintain their dynamic functions.

### Cellular importance of regulating stem loop structures with exposed GC-rich CSs

Interestingly, the defects in gene expression induced by stable stem loops and exposed GC-rich CSs in engineered *nos* mRNAs compared to WT *nos* are not mirrored in *osk*, *bcd*, and *HIV* RNAs (*38–44*). One possible explanation is that germ granules may lack protectors that recognize these motifs, causing the cells to identify these engineered mRNAs as abnormal and trigger downstream effects. *Drosophila* Staufen is an important regulator for *osk* mRNA localization and translation (*76*), as well as for the localization of *bcd* mRNA (*77*). The exposed GC-rich CS within a stem loop on *Drosophila osk* mRNA 3′UTRs is predicted to be a binding site for Staufen (*78, 79*). Similarly, the exposed GC-rich CS within a stem loop on *bcd* 3′UTR overlaps with Staufen binding motif (*77*). Therefore, these structural motifs not only enhance intermolecular mRNA base pairing but, equally important, serve as scaffolds to recruit RNA binding proteins essential for the post-transcriptional regulation of the mRNAs.

In the case of *HIV*, the SL1 is bound by the nucleocapsid domain of the Gag protein, which enhances genome dimerization for packaging *in vivo* (*77*). In addition to stabilizing intermolecular base pairing between *HIV* genomic RNAs, the Gag protein can multimerize to encapsulate the genome into virions, protecting *HIV* genomes from host degradation and promoting the viral infectivity (*80*). However, it is feasible that in the absence of Gag protein, host mRNAs modified with HIV-derived stem loops and exposed GC-rich CSs could trigger a host immune response. While this response may not trigger a global cellular response, it may be sufficient to specifically downregulate the gene expression of the modified host mRNAs.

Alternatively, these phenotypes could be driven by mRNA decay triggered by an elevated GC content and strong stem loops located in the 3′UTR (*66, 81*). Indeed, the HIV-derived SL1 structure has a GC-content of 71%, which is more than two-fold higher than the GC content of the average 3′UTR in *Drosophila* (*82*) including *nos* (32%). Therefore, a similar GC-content dependent mRNA decay might target *nos-non*, *nos-HIV* and *nos-nos* mRNAs generated in this study.

It is also unclear how exposed GC-rich CSs within stable stem loops exacerbate the embryonic and oogenesis phenotypes we observed (Figure 5,6). In embryos, only germ granule-localized *nos* translates (*53, 54, 56, 83*) and Nanos protein produced is required for abdominal patterning of the embryo. Therefore, as shown previously, the efficiency of abdominal segmentations and egg hatching correlates with Nanos protein levels produced at the posterior pole (*68, 69*). Notably, however, while the three CRISPR lines produced the same amount of Nanos protein, the translation of *nos-HIV* and *nos-nos* mRNAs is not normal. Specifically, the two *nos-HIV* and *nos-nos* localize 1.9 to 2.3-fold more mRNA than *nos-non* (Figure 4i), respectively, indicating that they should produce 1.9 to 2.3-fold more Nanos protein than *nos-non*, respectively. Regardless, this small difference in translational interference is not causing the exacerbated developmental phenotypes associated with exposed GC-sequences as the total amount of Nanos protein was similar among *nos-non*, *nos-HIV* and *nos-nos* (Figure 4k, j). Instead, the GC-rich sequences likely interfere with the downstream functions of Nanos protein. It is conceivable that the GC-rich CSs generate stable interactions and structural motifs not normally present in WT *nos* 3′UTR, which could be recognized by quality control regulators that hinder the proper functions of Nanos coded by an engineered, “foreign” mRNA. Which quality control mechanisms are at play and how they interfere with Nanos protein function is unclear.

RNA clustering has been observed in germ granules in *C. elegans* and zebrafish (*3, 84*) as well as P-bodies in *C. elegans* (*85*). Given that the core functional and structural principles are shared among these RNA granules (*1, 8*), our findings suggest that RNAs in these granules might be similarly compacted. This compaction could provide a mechanism by which the folding of mRNA controls intermolecular base pairing among diverse granule mRNAs. Interestingly, mRNAs become compacted upon localization to stress granules (*86, 87*). Therefore, along with RNA helicases (*19*), this structural compaction could further regulate intermolecular base pairing within stress granules and protect mRNA functional integrity for the time when cellular stress is over and when stress granule mRNAs return to their normal cellular functions. Collectively, our findings highlight the role of RNA structure in safeguarding mRNAs across cellular environments as well as the role of proteins in organizing mRNAs in cells.

## ACKNOWLEDGEMENTS

We would like to thank Drs. Anthony Leung and Sarah Woodson for the critical reading of our manuscript. We would like to thank Dr. Akira Nakamura for sharing the Nanos protein antibody with us. ST wrote the initial draft of the manuscript with all the authors participating in the review and editing of the manuscript. ST and TT: conceptualization. ST, HN, ZY: investigation and analysis. DT, HN: *In silico* modeling/analysis of monomers and homodimers. SR: Training with DMS-MaPseq technology. DT’s research was supported by grants CHE 2320256 and F-0019 from the National Science Foundation and the Welch Foundation, respectively. This research was supported by the NIGMS R35GM142737 grant awarded to TT.

## METHODS

### EXPERIMENTAL MODEL AND SUBJECT DETAILS

#### Fly stocks

Fly stocks and crosses were maintained at 25℃ on standard cornmeal/agar media. To test the base pairing capacity of mRNAs *in vivo*, we first crossed flies that expressed two *nos*-chimeras. The first chimera termed antisense *nos* 3′UTR, expressed the endogenous *nos* gene fused with the 3′UTR composed of the +2 localization element derived from the *nos* 3′UTR (*83*) and antisense sequence of *nos* 3′UTR (region spanning nucleotides 658 to 159). This chimera was generated by initially inserting two ATTP sites into the last intron of the *nos* gene using CRISPR/Cas9 genome editing, which allowed subsequent site-specific recombination of ATTB-flanked DNA *via* PhiC31 recombination by Fungene, as described previously (*88*). This approach enabled the reconstitution of the *nos* CDS and the simultaneous replacement of its 3′UTR with an RNA sequence of choice. The second chimera was transgenically expressed using a reporter mRNA fused with a CDS of a far-red fluorescent protein (IRFP670) and the 3′UTR of WT *nos* (*36*). Its expression was driven by the Gal4-responsive promoter. To generate flies expressing *nos* with exposed *non*, *HIV*, and *nos* complementary sequences, we inserted exposed *non*/*HIV/nos* palindromes in the *nos* genome location (*nos-non, nos-HIV,* and *nos-nos*, respectively), using the CRISPR/Cas9-PhiC31 approach described above. After eight rounds of background crossing with balancer flies and subsequent removal of balancer chromosomes, the flies expressing these *nos* gene variants were crossed with *nos^def^* (*73*) or *nos^BN^* flies (*65*) to study the effects of *nos-non, nos-HIV,* and *nos-nos* on fly oogenesis and embryogenesis, respectively.

#### *Drosophila* cell culturing

*Drosophila* Ras-attP-L1 cell line (DGRC Stock 249) was maintained at 25℃ as described previously (*89*). Schneider’s insect medium (Sigma: S0146) supplemented with 10% FBS (Thermo Fisher: 26140079) and 1X penicillin-streptomycin (Thermo Fisher: 15070063) was used to maintain the cell line.

## METHOD DETAILS

### *In vitro* RNA transcription

DNA templates containing the T7 promoter for *in vitro* transcription were ordered from IDT and amplified by Q5 high-fidelity DNA polymerase (NEB: M0492L). The amplified DNA was purified with Zymo DNA clean & concentrator kit (Zymo: D4003T). For *in vitro* RNA clustering assays, transcription templates ranging from 200-500 ng were transcribed *in vitro* using the MEGAscript T7 transcription Kit (Thermo Fisher: AM1333). 20 μL of transcription reaction was carried out for 6 hours (hrs) at 37℃. Following the transcription reaction, each reaction mixture was treated with 1 μL of TURBO DNase (Thermo Fisher: AM2238) for 15 mins at 37℃. The reaction was stopped with ammonium acetate stop solution following the manufacturer’s instructions. Then the *in vitro* transcribed RNAs were purified using phenol:chloroform extraction (Thermo Fisher: 15593031) following the manufacturer’s protocol. The purified RNAs were stored in isopropanol (Thermo Fisher: 278475-1L) at -20℃ for a maximum of one month to prevent degradation.

### *In vitro* intermolecular RNA base pairing assays and gel electrophoresis

The protocol was adapted from (*39, 41*). *In vitro* transcribed RNAs suspended in isopropanol were centrifuged at 14,000 g for 15 mins at 4℃ to precipitate the RNAs. The RNA pellet was washed twice with 80% RNase-free ethanol (Thermo Fisher: BP2818500) by centrifugation at 14,000 g for 2 mins at 4℃. The RNA pellet was then air-dried and resuspended in 20 μL of nuclease-free water (Thermo Fisher: AM9937) in a 1.5 mL test tube on ice. The concentration and quality of the RNA were accessed using a NanoDrop spectrometer.

RNA samples resuspended in water were denatured at 90℃ for 2 mins, and afterward the sample tubes were immediately placed on the ice for at least 15 mins. For a reaction using one RNA sample, 32 pmol of denatured RNA was aliquoted into a 200 μL PCR tube and mixed with 2 μL of 5X RNA gel refolding buffer (50mM sodium cacodylate (pH 7.5) (Electron Microscopy Science: 11654), 300mM KCl (Sigma: 60128-250G-F) and 5mM MgCl_2_ (Sigma: M1028-100ML) and nuclease-free water added to the final volume of 10 μL. The mixture was refolded at RT for 1.5 hrs. For a rection using two RNA species, 16 pmol of each RNA sample was mixed with 1 μL of 5X RNA gel refolding buffer and nuclease-free water to reach a final volume of 5 μL, and was refolded separately at RT for 1 hr. Then the two refolded RNA samples were combined and incubated for an additional 30 mins at RT. A 2% agarose gel supplemented with 1X SYBR safe (Thermo Fisher: S33102) and gel running buffer (0.5× TAE (Thermo Fisher: 15558042), 0.1mM MgCl_2_) was prepared and prechilled at 4℃. To each refolding reaction, 1.9 μL of formamide-free loading dye (Thermo Fisher: R0611) was added. After loading all samples and the RNA ladders (NEB: N0362S) onto the gel, the gel was run for 2 hrs at 70V and at 4℃ to separate RNA populations. The gel was imaged using a ChemiDoc imaging system. The list of RNA sequences used for RNA gel electrophoresis is reported in Table S1.

### RNA clustering reaction

This protocol was modified based on (*20, 28*). On the day of the experiment, a 10X RNA cluster refolding buffer (200 mM KCl, 100 mM MgCl_2,_ and 100 mM Tris (pH 7.0) (Millipore: 648314-100ML)) was freshly prepared and stored at RT. A 1 mL aliquot of filtered 100 mM spermine-tetrahydrochloride (Sigma: S1141-1G) in nuclease-free water and 50% PEG8000 (NEB: B1004SVIAL) were thawed on ice. Note that the 50% PEG8000 was used within 2 months of opening to ensure consistent results due to potential degradation.

The preparation of *shu-top* and *shu-bottom* RNA samples was similar to the one used for the *in vitro* intermolecular RNA base pairing assays and gel electrophoresis. However, the *in vitro* transcription reaction contained Cy5-labeled UTPs (aminoallyl-UTP-Cy5) (Jena Bioscience: NU-821-CY5). The concentration of the labeled UTP was adjusted to ensure an average of 3 labeled uracils per RNA.

For the separate folding condition, 32 pmol of each *shu-top* and *shu-bottom* RNAs were separately denatured at 90℃ for 2 mins and the sample tubes were immediately placed on ice for 15 mins. For each RNA sample, 2 μL of the 10X RNA cluster refolding buffer and nuclease-free water were added to the RNAs to reach a final volume of 7 μL for each RNA sample. The folding reaction was incubated at RT for 1 hr. Afterward, the folded *shu-top* and *shu-bottom* RNAs were combined. 6 μL of 1:2 (vol:vol) 100 mM spermine and 50% PEG8000 premix were added and thoroughly mixed due to the stickiness of PEG8000, resulting in a final volume of 20 μL with each RNA 1.6 μM per reaction.

For the co-folding condition, 32 pmol of each *shu-top* and *shu-bottom* RNAs were denatured at 90℃ for 2 mins together and the sample tube was immediately placed on ice for 15 mins. The clustering reaction, which had a final volume of 20 μL with each RNA 1.6 μM per reaction, was the same as the one for separate folding. The sequences of *top* and *bottom* RNAs are listed in Table S1.

### Fluorescence assays to detect intermolecular base pairings in *in vitro* RNA clusters

To prepare the staining solution, 780 μL of nuclease-free water was added to 5mg lyophilized DFHBI-1T (LUCERNA: 410-5MG), resulting in a stock DFHBI-1T staining solution with a concentration of 20 mM. The solution was stored at -20℃. Next, 2 μL of 2 mM DFHBI-1T was mixed into the final 20 μL clustering reaction to reach 0.1 mM working concentration, and the reaction was incubated in the dark for 4 hrs at RT. For co-folding and separate folding, the ratio of DFHBI-1T (0.1 mM): *shu-top* (1.6 μM): *shu-bottom* (1.6 μM) is 62.5:1:1. Because a base-paired *top* and *bottom* can form a broccoli structure with two intercalated DFHBI-1T molecules (*48*), we estimate an approximate 30-fold excess of DFHBI-1T to the potential broccoli structures in our experiments. Afterwards, 8-10 clusters were randomly selected for imaging using VT-iSIM with a 100X 1.5 NA oil immersion objection and a z-series of 27 slices with a step size of 150nm. Two experimental replicates were performed for each condition and time point.

To detect intermolecular base pairing by RNA gel electrophoresis, 16 pmol of each Cy5 (Jena Bioscience: NU-821-Cy5) labeled *shu-top* and *shu-bottom* RNAs were used following the protocol described for *in vitro* intermolecular RNA base pairing assays and gel electrophoresis. After the electrophoresis was complete, the gel was stained with 5 μM DFHBI-1T at RT for 15 mins and then imaged using Amersham™ Typhoon™ 5 scanner (GE Healthcare) with 488 and Cy5 filters as described before (*45*).

### Embryo collection and germ granule isolation

Embryos were collected as described previously (*90*). The granule isolation was adapted from (*51*). Approximately fifty caged flies were allowed to lay eggs at 25℃ on a fresh apple juice agar plate supplemented with yeast paste for 1.5 hrs. About 20 μL of embryos were collected in 1X PBS in a 1.5mL test tube. After the embryos settled at the tube bottom, the 1X PBS was replaced with 150 μL of freshly made 1X cold lysis buffer (0.34M sodium cacodylate (pH 7.5), 6mM MgCl_2_, 1X protease inhibitor complete mini EDTA (Millipore: 11836170001) free and 1U/ μL RNase inhibitor (Thermo Fisher: 10777019) adapted from (*91*). The embryos were then lysed in lysis buffer in the presence of 20 μL of 0.1mm glass beads (Millipore: G1145-10G) using a cordless homogenizer for 2 mins at RT. The lysate was clarified by centrifugation at 2000 g for 2 mins and the supernatant, which was separated from the debris and the beads, was transferred to a new test tube. The supernatant was then centrifuged again at 10,000 g for 15 mins at 4℃, and about 120 μL of the soluble fraction without touching the bottom pellet was transferred to a new test tube. This soluble fraction was re-centrifuged at 10,000 g for 15 mins at 4℃. About 100 μL of the final clarified soluble fraction, which represented the fraction outside the germ granules, was transferred to a new test tube. The pellet in the initial sample tube, which represented the germ granule fraction, was washed three more times with 100 μL of 1X cold lysis buffer and centrifuged at 10,000 g for 5 mins each time at 4℃. Finally, the pellet was suspended in 100 μL of 1X cold lysis buffer. Both fractions were temporally stored on ice before proceeding to the DMS treatment.

### DMS treatment and total RNA isolation

The experimental procedures were adapted from (*49, 91*). Before DMS treatment, the sample tubes containing the isolated germ granule fraction and the fraction outside the germ granules were incubated at 26℃ for 10 min. Then 10 μL of 100% DMS (Millipore: D186309-5ML) or nuclease-free water as a negative control was added to both fractions. The samples were incubated on a thermomixer at 800 rpm, 26℃ for 5 mins, and then immediately put on the ice with the addition of 60 μL 100% 2-mercaptoethanol (Millipore: 444203) to stop the reaction. Next, 500 μL of TRIzol (Thermo Fisher: 15596026) was added to the samples, and total RNA was purified using the Zymo direct-zol RNA miniprep kit (Zymo: R2050). The purified RNA was eluted in water and stored at -80℃. For each sample replicate, at least 7 rounds of egg collection and granule isolation were performed. The total RNA samples were pooled and concentrated using the Zymo RNA clean & concentrator kit (Zymo: R1014) to obtain sufficient RNA material. The RNA samples were then treated with TURBO DNase and reversed transcribed using gene-specific reverse primers, as described previously (*49*). The regions of interest for DMS structural probing, with a size of approximately 200 nts, were PCR amplified using Q5 high-fidelity polymerase. The resulting DNA fragments were extracted using SizeSelect 2% precast gel (Thermo Fisher: G661012) on the E-Gel Power Snap Electrophoresis Device and stored at -20℃. The primer sequences used in DMS-MaPseq are listed in Table S2.

### DNA library preparation and sequencing

The concentration of each extracted DNA fragment was determined using Qubit 4 fluorometer with 1X dsDNA HS assay kit (Thermo Fisher: Q32854). For each library preparation, 100 ng of pooled DNA fragments were used as input and prepared using the NEBNext Ultra II DNA library prep kit for Illumina (NEB: E7645S) and its protocol. The prepared DNA libraries were sent for sequencing using the MiSeq system by the Johns Hopkins Genetic Core Facility.

### RNA isolation and qRT-PCR

About fifty caged flies were allowed to lay eggs at 25℃ on a fresh apple juice agar plate with a scope of yeast paste for 1.5 hrs. The embryos were collected, homogenized in Trizol, and stored at -80℃ until the next step. Total RNA was extracted using chloroform (Thermo Fisher: C298-500) and precipitated in isopropanol following the TRIzol Reagent User Guide. Next, about 2 μg of total RNA was treated with RQ1 RNase-Free DNase (Promega: M6101) following the manufacturer’s instructions. cDNA synthesis was performed using SuperScript^TM^ III reverse transcriptase (Thermo Fisher: 18080093) following the manufacturer’s protocol. For each 10 μL of qRT-PCR reaction, approximately 100 ng of cDNA, 1.5 μL of 1 μM forward and reverse primers, and 5 μL iTaq universal SYBR® Green supermix (Bio-Rad: 1725122) were added. Quantitative PCR analysis was performed using a CFX Opus 96 Real-Time PCR System from Bio-Rad. The primer sequences are listed in Tables S2 and S4.

### Single-molecule fluorescent in situ hybridization (smFISH) on fly embryos

The embryo collection was carried out as described before (*90*). The embryos were stored in 100% methanol at 4°C until the smFISH experiments. To label individual mRNAs, commercially available Stellaris probes were used. Each set of probes consisted of 30 to 48, 20-nt DNA oligos designed with the default setting on Stellaris Probe Designer. Each oligo was covalently conjugated with either CAL Fluor 590 or Quasar 670. In-house labeling of smFISH probes involved designing the probe set with Stellaris Probe Designer and ordering the oligos from IDT. Subsequently, the oligos were modified using terminal deoxynucleotidyl transferase (Thermo Fisher: EP0161) and amino-11-ddUTP (Lumiprobe: 15040), followed by covalent conjugation with AF488 (Lumiprobe: 21820), AF568 (Lumiprobe: 24820) or AF647 (Lumiprobe: 26820) NHS esters as previously described (*92*). Hybridization of mRNAs with smFISH was performed as previously described (*90*). The sequences of smFISH probes are listed in Table S3 or adapted from previous (*17, 36*).

### Simulation of RNA structure

We performed all the simulations using an updated version of the Single Interaction Site (SIS) model (*34*). The modification restricts the number of multiple base pairs a single nucleotide can form simultaneously. The simulations were performed on Graphics Processing Units (GPUs) using a custom OpenMM code (*93*) to enhance sampling of the conformational space. We used low-friction Langevin dynamics, in which the viscosity of water was reduced by a factor of 100 (*94*). Even for the SIS model for RNA, the simulations are computationally extensive, thus requiring the simulated tempering method to ensure that the conformational space is sampled exhaustively (*95*). The trajectories were analyzed using the Multistate Bennett Acceptance Ratio (MBAR) to calculate all properties of interest (*96*). All simulations were performed with 1M NaCl, where electrostatic interactions are weak, and only base pairing interactions dominate (*97*).

#### Monomer simulations

A single RNA molecule in the extended conformation was initially placed in a simulation box (the size of the box is much larger than the RNA size and does not play any role in determining RNA structures. Simulations were performed for 5 x 10^9^ time steps, in which the first 5 x 10^8^ steps were discarded, which ensures that only equilibrated structures are used in computing various quantities of interest.

#### Homodimer simulations

Two representative snapshots from monomer simulations were randomly picked and placed in a sphere of radius *R*. The two RNAs were constrained inside the sphere by defining the following potential for any particles:

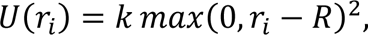

where *r*_*i*_ is the distance between the particle *i* and the sphere center. *R* was chosen big enough so that the two RNAs have enough space to adopt extended conformations, but also small enough to prevent the two chains from drifting away from each other. Simulations were then conducted for 5 × 10^9^time steps.

#### Base accessibility calculations

Base accessibility of the nucleotide *i* is defined as *H*_*i*_ = *S*_*i*_γ_*i*_, where *S*_*i*_ is the solvent-accessible surface area of nucleotide *i* (calculated using the Lee-Richards algorithm (*98*) in FreeSASA (*99*). γ_*i*_ adopts two values: 0-if the nucleotide *i* is involved in base pairing and 1-otherwise. We surmise that the base accessibility is roughly proportional to the DMS-MaPseq output from experiments. In addition to the RNA secondary structure (reflected in γ_*i*_), the accessibility of a base also depends on the RNA tertiary structure (reflected in *S*_*i*_). If a base is deeply buried in the core of the RNA, its accessibility is small. On the other hand, if the base is located near the RNA periphery, its likelihood to form intermolecular interactions is higher.

#### Clustering of secondary structures

RNA conformations are grouped based on the secondary structures as following. Each conformation *i* is fully specified by a set of base pairs *B*_*i*_ = {{*a, b*},…}, where *a* and *b* are the two nucleotides that base pair to each other. The similarity between the two conformations *i* and *j* are calculated using the Jaccard distance between the two sets *B*_*i*_ and *B*_*j*_ as:

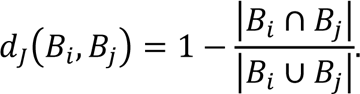

The Jaccard distance considers the difference in sizes of two sets and is bounded between 0 and 1. Thus, if the two conformations share no base pair, then *d*_*J*_(*B*_*i*_, *B*_*j*_) = 1, whereas *d*_*J*_(*B*_*i*_, *B*_*j*_) = 0 if *B*_*i*_ = *B*_*j*_.

We then generated the distance matrix *J*, where *J*_*ij*_ = *d*_*J*_(*B*_*i*_, *B*_*j*_) for the pair *i* and *j*. *J* was then used as the input to HDBSCAN (*100*), which is a density-based clustering algorithm, to extract clusters of RNA conformations. The free energies of the clusters were subsequently calculated using MBAR.

### Transfection for *Drosophila* Ras cells

*Drosophila* Ras cells were seeded on the chambered cell culture slides (Grace Bio-Labs: 103510) on the day of transfection. 200 ng of plasmids containing LacZA and LacZB sequences, each tagged with *HIV, nos, non, sense* or *antisense* CSs and fused with metallothionein promoters (*101*), were transfected into the cells using effectene transfection reagent (Qiagen: 301425). After 24 hrs, 0.5 mM copper sulfate (Sigma: C8027-500G) was added to the cell culture to induce the expression of *LacZA* and *LacZB* RNAs. After 24 hrs of induction, the cells were fixed with 4% formaldehyde for 10 mins and subsequently washed with 1X PBS. To detect *LacZA* and *LacZB* RNAs, HCR™ commercially designed probes and RNA-FISH protocol (Molecular instruments) for adherent cells were used following the manufacturer’s instructions.

### Egg hatching assays

Approximately 20 *non/ nos^BN^*, *HIV/ nos^BN^* or *nos/ nos^BN^* virgin females were crossed with 10 WT (*W^1118^*) young males. The crosses were maintained at 25℃ for 3 days on standard cornmeal/agar media supplemented with yeast powder. Then the flies were caged and supplied with a fresh apple juice plate containing a dollop of yeast paste for 24 hrs at 25℃ and allowed to lay eggs. Afterward, the plate was collected and the number of eggs on it was counted and replaced with a new apple juice plate. The plate was then kept at 25℃ for an additional 24 hrs to count the number of hatched eggs. The hatching rate was determined by dividing the number of hatched eggs by the total number of eggs on the same plate.

### Western Blot analysis

The embryos were collected and flash-frozen using liquid nitrogen. The samples were stored at - 80℃ until the next step. The samples were homogenized in 100 μL of cold lysis buffer (150 mM sodium chloride, 50 mM pH 8.0 Tris-HCL, 1% Triton-X100, 0.5% sodium deoxycholate, and 0.1% SDS). After incubating on ice for 20 mins, the lysates were centrifuged at 15,000 g for 20 mins at 4℃. The resulting supernatants were transferred to new 1.5 mL tubes, and the protein concentrations were quantified using the Pierce^TM^ BCA protein assay kit (Thermo Fisher: 23227) and Nanodrop, following the manufacturer’s protocol. About 20-50 μg of total proteins were mixed with 1X Laemmli (Bio-Rad: 1610747) and 50 mM DTT (Thermo Fisher: R0861), resulting in a total final volume of 30 µL. The samples were then boiled at 95℃ for 5 mins. Once cooled, they were loaded into a 7.5% Criterion™ TGX Stain-Free™ Protein Gel (Bio-Rad: 5671024). 10µL Precision Plus Protein™ WesternC™ blotting standards (Bio-Rad: 1610376) was used as ladders. The gel was run in 1X Tris-glycine SDS running buffer (2.5 mM Tris, 19.2 mM glycine (Sigma: G8898-500G), 0.01% SDS (Thermo Fisher: 28364)) at 150 V for 1.5 hrs at 4℃. After electrophoresis, the bands were transferred to a Trans-Blot Turbo Midi 0.2μm Nitrocellulose membrane (Bio-Rad: 1704159) using Trans-Blot Turbo transfer system and Bio-Rad mixed molecular weight protocol. The membrane was then blocked in 5% blotting grade non-fat dry Milk (Bio-Rad: 1706404) in PBST (1x PBS (Thermo Fisher: 70011044), 0.1% Tween-20 (Sigma: 655204)) for 1 hr at RT on a rocker. Next, the membrane was incubated overnight at 4℃ with primary, mouse anti-β-actin antibody (Abcam: ab8224), which was diluted in 1% blotting grade non-fat dry milk/PBST to 1:2000. After incubation, the membrane was washed three times with PBST and incubated with secondary, goat anti-mouse IgG (HRP) antibody (Abcam: ab6789), which was diluted in 1% blotting grade non-fat dry milk/PBST to 1:10000 for 2 hrs. The membrane was washed five times with PBST. The signal was developed using SuperSignal^TM^ western dura extended duration substrate (Thermo Fisher: 34075). The membrane was imaged by ChemiDoc MP with an exposure time of 10-60 ms. To detect Nanos protein expression on the same membrane, the previous primary and secondary antibodies were removed by western blot stripping buffer (Thermo Fisher: 21059) following the manufacturer’s instructions. Rabbit anti-Nanos antibody (a gift from Nakamura Lab) (1:1000) (*102*), and goat anti-rabbit IgG (HRP) antibody (Abcam: ab6721) (1:10000), were then used to detect Nanos following the protocol described for detecting β-actin. Finally, for each protein, two to three biological replicates per sample were run on the same western blot. The analysis was performed following the guidelines from ImageJ User Guide-30.13 Gels.

### Immunostaining of fly ovaries

The females were fed with yeast powder the day before the dissection. The ovaries were dissected in Schneider’s insect medium supplied with 200 μg/mL insulin (Sigma: I5500-500MG) at RT. The tissues were fixed in 4% formaldehyde (Electron Microscopy Sciences: 15713) in PBTx (1XPBS, 0.1% Triton-X100 (Millipore: TX1568-1) for 20 mins at RT and then washed twice in PBTx, each time for 10 min. Subsequently, the tissues were blocked overnight at 4℃ in BBTx (1XPBTx, 0.5% BSA (Millipore: A3294-50G), 2% NGS (Abcam: ab7481)). Rabbit anti-Vasa antibody (RRID: AB2940894) (*51*) and mouse anti-1B1 antibody (DSHB: AB528070), were diluted in BBTx to 1:500 and 1:50, respectively. They were added to tissues and incubated overnight at 4℃. Afterward, the tissues were washed twice in PBTx for 10 mins at RT and then treated with goat anti-rabbit IgG Alexa488 antibody (Thermo Fisher: A-11070) and goat anti-mouse IgG Alexa568 antibody (Thermo Fisher: A-11004), which were diluted in BBTx to 1:1000, for 4 hrs at RT. The tissues were then washed twice in PBTx for 10 mins at RT. To stain the DNA, 1 μg/mL DAPI (Sigma: 10236276001) diluted in PBTx was added to the samples and incubated for 5 mins at RT. Finally, the samples were mounted on slides using ProLong™ glass antifade mountant (Thermo Fisher: P36980) and cured overnight at RT before imaging.

### Microscopy and deconvolution

Images were acquired with a vt-instant Structured Illumination Microscope (vt-iSIM; BioVision Technologies) equipped with the 405nm 100mW, 488nm 150mW, 561nm 150mW, 642nm 100mW, and 445nm 75 mW lasers, two ORCA-Fusion sCMOS cameras and the Leica HC PL APO 63x/1.30 GLYC CORR CS2, HC PL APO 63x/1.40 OIL CS2 and HC PL APO 100x/1.47 OIL CORR TIRF objectives as described before (reference 26). Images were acquired in three dimensions (3D) and then deconvolved using Huygens (Scientific Volume Imaging).

### Identification of CSs

The identification of CSs involves scanning through all possible pairs of starting positions, considering the forward direction of sequence 1 and the reverse direction of sequence 2. Sequences 1 and 2 were identical in order to identify the CSs which were self-complementary. Starting with each possible position in sequence 1, the program checks for complementary nucleotides in the reverse direction of sequence 2. If a match is found between a nucleotide in sequence 1 and its corresponding nucleotide in sequence 2, the program extends the length of the complementary sub-sequence by one, continuing the process for subsequent nucleotides. This extension continues until a mismatch occurs, at which point the program records the CS if the length and/or the GC content passes the threshold. The program then moves on to the next pair of starting positions and repeats the process until all possible positions are evaluated. The identified CSs on 3′UTRs of *nos*, *pgc* and *gcl* are listed in Tables S5-S7, respectively.

Codes are available Github: https://github.com/AnneyYeZiqing/sticky_finder.

## QUANTIFICATION AND STATISTICAL ANALYSIS

### Quantifying the level of fold enrichment in *in vitro* RNA clusters

The images were imported into FIJI/Image J (*103*), and the channels of DFHBI-1T and RNA were split. A 5×5 pixel rectangle was used to measure the levels of the integrated density inside and outside the assemblies. For each image, three different regions were randomly selected using a 5×5 pixel rectangle. The levels of the integrated density inside and outside the clusters were determined in the RNA channel. The fold enrichment was calculated as Integrated Density _(inside)_/ Integrated Density _(outside)._

### Quantifying the level of intermolecular base pairing in *in vitro* RNA clusters

Same as above, three different regions inside and outside the assemblies were randomly chosen and measured. The background signal was determined by the dye-only condition without the addition of RNAs. The normalized integrated density of the selected region was calculated as the Integrated Density of the dye _(selected region)_-Integrated Density _(dye background)_/Integrated Density of RNA _(selected region)_.

### Quantifying mRNA fold enrichment in isolated germ granules

Germ granules were isolated as described above and previously (26). However, from the last washing step, 100 μL of the soluble fraction in 1X cold lysis buffer was collected and the pellet was suspended in 100 μL 1X cold lysis buffer. The total RNA from these two fractions was extracted by Zymo Direct-zol RNA Miniprep kit and used for cDNA synthesis as described previously (see Methods Details: RNA isolation and qRT-PCR). For each qRT-PCR reaction of each sample, 100±10ng of cDNA was used as input. Two biological replicates with three technical replicates were analyzed. To calculate the fold enrichment of a particular gene (here termed X), _2_−(Cq(gene X in soluble)– Average of Cq(gene X in soluble)) was used for calculating soluble fractions. 2−(Cq(gene X in pellets)– Average of Cq(gene X in soluble)) was used for pellets. Primers are listed in Table S2.

### Analysis of DMS-MaPseq data

The raw fasta files were analyzed using the **D**etection of **R**NA folding **E**nsembles using **E**xpectation-**M**aximization (DREEM) as previously described (*63*). The code for this analysis can be found at https://codeocean.com/capsule/0380995/tree. Two biological replicates were used. For DMS-MaPseq outside the germ granules, Pearson correlation coefficients (r) are 0.93, 0.97 and 0.99 between the replicates of *nos, pgc* and *gcl* 3′UTRs, respectively. For DMS-MaPseq within the germ granules, r = 0.99, 0.98 and 0.89 in *nos, pgc* and *gcl* 3′UTRs, respectively.

Averages of the two replicates were taken for subsequent analysis. The profiles of average reactivity (K=1 in DREEM) were used to determine the structuredness of CSs using RNA probing which was set to use Diegan et al. 2009 (*104*) and temperature at 26℃. Predicted secondary structures were visualized by FORNA (*105*) and VARNA (*106*). To compare the similarity in DMS profiles between inside and outside the germ granules, the reactivity of each informative nucleotide for each RNA sequence was correlated between the inside and outside of the germ granules. Pearson’s Correlation coefficient was then calculated. Additionally, to compare changes in the DMS reactivity of each nucleotide between inside and outside the germ granules, the DMS reactivity of each nucleotide inside the germ granules was divided by the corresponding value outside the granules. The significant thresholds were calculated based on the variance in DMS reactivity between two biological replicates under the same condition.

### Quantifying the co-localization of germ granule mRNA assemblies by PCC(Costes) and PCC

Analyses were performed using PCC(Costes) and PCC co-localization ImageJ plugin (*107*) as described before (*36*).

### qRT-PCR analysis of transcripts in *nos-non, nos-HIV* and *nos-nos* flies

*Drosophila GADPH* was used as a control to calculate the relative transcript levels, which was 2^−(Cq(region of interest)– Average of Cq(*GAPDH*))^. Then the data were further normalized to *nos-non*. Primers are listed in Table S4.

### Quantifying mRNA concentration and the cluster abundance in embryos

Analyses were performed using Aircalize spot detection algorithm (*108*) as described before (*36*).

### Western blot analysis

For each protein target, different samples with two to three biological replicates were run on the same western blot. The analysis was performed following the guidelines from ImageJ User Guide-30.13 Gels.

### Counting female GSCs

Female GSCs were counted based on the morphology of the anti-1B1 staining and juxtaposition to the terminal filament as described previously (*109*).

## SUPPLEMENTAL INFORMATION

**Table S1. RNAs used for in vitro experiments, related to Figures 1, S1, S5, S6 and STAR methods.**

**Table S2. Primers used in DMS-MaPseq on the isolated granules, related to Figures 2, S1, S2 and STAR methods.**

**Table S3. smFISH probes, related to Figure 2 and STAR methods.**

**Table S4. Primers used in *nos* qRT-PCR and PCR, related to Figure 4 and STAR methods.**

**Table S5. CSs on *nos* 3’UTR, related to Figure 7**.

**Table S6. CSs on *pgc* 3’UTR, related to Figure 7**.

**Table S7. CSs on *gcl* 3’UTR, related to Figure 7**.

## REFERENCES

1. S. Tian, H. A. Curnutte, T. Trcek, RNA Granules: A View from the RNA Perspective. Molecules 25, (2020).

2. S. F. Banani, H. O. Lee, A. A. Hyman, M. K. Rosen, Biomolecular condensates: organizers of cellular biochemistry. Nat Rev Mol Cell Biol 18, 285–298 (2017).

3. C. Eno, C. L. Hansen, F. Pelegri, Aggregation, segregation, and dispersal of homotypic germ plasm RNPs in the early zebrafish embryo. Dev Dyn 248, 306–318 (2019).

4. D. S. W. Protter, R. Parker, Principles and Properties of Stress Granules. Trends Cell Biol 26, 668–679 (2016).

5. C. L. Riggs, N. Kedersha, P. Ivanov, P. Anderson, Mammalian stress granules and P bodies at a glance. J Cell Sci 133, (2020).

6. S. M. Sagan, S. C. Weber, Let’s phase it: viruses are master architects of biomolecular condensates. Trends Biochem Sci 48, 229–243 (2023).

7. E. Voronina, G. Seydoux, P. Sassone-Corsi, I. Nagamori, RNA granules in germ cells. Cold Spring Harb Perspect Biol 3, (2011).

8. A. Chiappetta, J. Liao, S. Tian, T. Trcek, Structural and functional organization of germ plasm condensates. Biochem J 479, 2477–2495 (2022).

9. P. Yang et al., G3BP1 Is a Tunable Switch that Triggers Phase Separation to Assemble Stress Granules. Cell 181, 325–345 e328 (2020).

10. J. Smith et al., Spatial patterning of P granules by RNA-induced phase separation of the intrinsically-disordered protein MEG-3. Elife 5, (2016).

11. M. Bose, M. Lampe, J. Mahamid, A. Ephrussi, Liquid-to-solid phase transition of oskar ribonucleoprotein granules is essential for their function in Drosophila embryonic development. Cell 185, 1308–1324 e1323 (2022).

12. B. A. Chau, V. Chen, A. W. Cochrane, L. J. Parent, A. J. Mouland, Liquid-liquid phase separation of nucleocapsid proteins during SARS-CoV-2 and HIV-1 replication. Cell Rep 42, 111968 (2023).

13. A. Patel et al., A Liquid-to-Solid Phase Transition of the ALS Protein FUS Accelerated by Disease Mutation. Cell 162, 1066–1077 (2015).

14. A. Wang et al., A single N-terminal phosphomimic disrupts TDP-43 polymerization, phase separation, and RNA splicing. EMBO J 37, (2018).

15. M. Feric et al., Coexisting Liquid Phases Underlie Nucleolar Subcompartments. Cell 165, 1686–1697 (2016).

16. A. Jain, R. D. Vale, RNA phase transitions in repeat expansion disorders. Nature 546, 243–247 (2017).

17. T. Trcek et al., Drosophila germ granules are structured and contain homotypic mRNA clusters. Nat Commun 6, 7962 (2015).

18. S. C. Little, K. S. Sinsimer, J. J. Lee, E. F. Wieschaus, E. R. Gavis, Independent and coordinate trafficking of single Drosophila germ plasm mRNAs. Nat Cell Biol 17, 558–568 (2015).

19. D. Tauber et al., Modulation of RNA Condensation by the DEAD-Box Protein eIF4A. Cell 180, 411–426 e416 (2020).

20. B. Van Treeck et al., RNA self-assembly contributes to stress granule formation and defining the stress granule transcriptome. Proc Natl Acad Sci U S A 115, 2734–2739 (2018).

21. R. Chen et al., HDX-MS finds that partial unfolding with sequential domain activation controls condensation of a cellular stress marker. Proc Natl Acad Sci U S A 121, e2321606121 (2024).

22. C. Sahin et al., Mass Spectrometry of RNA-Binding Proteins during Liquid-Liquid Phase Separation Reveals Distinct Assembly Mechanisms and Droplet Architectures. J Am Chem Soc 145, 10659–10668 (2023).

23. Clifford P. Brangwynne, P. Tompa, Rohit V. Pappu, Polymer physics of intracellular phase transitions. Nature Physics 11, 899–904 (2015).

24. K. M. Ruff et al., Sequence grammar underlying the unfolding and phase separation of globular proteins. Mol Cell 82, 3193–3208 e3198 (2022).

25. S. Basu et al., Rational optimization of a transcription factor activation domain inhibitor. Nat Struct Mol Biol 30, 1958–1969 (2023).

26. J. D. Schmit, M. Feric, M. Dundr, How Hierarchical Interactions Make Membraneless Organelles Tick Like Clockwork. Trends Biochem Sci 46, 525–534 (2021).

27. Y. Shin, C. P. Brangwynne, Liquid phase condensation in cell physiology and disease. Science 357, (2017).

28. E. M. Langdon et al., mRNA structure determines specificity of a polyQ-driven phase separation. Science 360, 922–927 (2018).

29. R. R. Poudyal, J. P. Sieg, B. Portz, C. D. Keating, P. C. Bevilacqua, RNA sequence and structure control assembly and function of RNA condensates. RNA 27, 1589–1601 (2021).

30. P. C. Bevilacqua, A. M. Williams, H. L. Chou, S. M. Assmann, RNA multimerization as an organizing force for liquid-liquid phase separation. RNA 28, 16–26 (2022).

31. J. W. Miller et al., Recruitment of human muscleblind proteins to (CUG)(n) expansions associated with myotonic dystrophy. EMBO J 19, 4439–4448 (2000).

32. A. R. Haeusler et al., C9orf72 nucleotide repeat structures initiate molecular cascades of disease. Nature 507, 195–200 (2014).

33. A. E. Renton et al., A hexanucleotide repeat expansion in C9ORF72 is the cause of chromosome 9p21-linked ALS-FTD. Neuron 72, 257–268 (2011).

34. H. T. Nguyen, N. Hori, D. Thirumalai, Condensates in RNA repeat sequences are heterogeneously organized and exhibit reptation dynamics. Nat Chem 14, 775–785 (2022).

35. T. Trcek, R. Lehmann, Germ granules in Drosophila. Traffic 20, 650–660 (2019).

36. T. Trcek et al., Sequence-Independent Self-Assembly of Germ Granule mRNAs into Homotypic Clusters. Mol Cell 78, 941–950 e912 (2020).

37. M. G. Niepielko, W. V. I. Eagle, E. R. Gavis, Stochastic Seeding Coupled with mRNA Self-Recruitment Generates Heterogeneous Drosophila Germ Granules. Curr Biol 28, 1872–1881 e1873 (2018).

38. H. Jambor, C. Brunel, A. Ephrussi, Dimerization of oskar 3’ UTRs promotes hitchhiking for RNA localization in the Drosophila oocyte. RNA 17, 2049–2057 (2011).

39. D. Ferrandon, I. Koch, E. Westhof, C. Nusslein-Volhard, RNA-RNA interaction is required for the formation of specific bicoid mRNA 3’ UTR-STAUFEN ribonucleoprotein particles. EMBO J 16, 1751–1758 (1997).

40. J. L. Clever, M. L. Wong, T. G. Parslow, Requirements for kissing-loop-mediated dimerization of human immunodeficiency virus RNA. J Virol 70, 5902–5908 (1996).

41. R. Marquet et al., Dimerization of human immunodeficiency virus (type 1) RNA: stimulation by cations and possible mechanism. Nucleic Acids Res 19, 2349–2357 (1991).

42. A. Mujeeb, J. L. Clever, T. M. Billeci, T. L. James, T. G. Parslow, Structure of the dimer initiation complex of HIV-1 genomic RNA. Nat Struct Biol 5, 432–436 (1998).

43. J. C. Paillart, E. Skripkin, B. Ehresmann, C. Ehresmann, R. Marquet, A loop-loop “kissing” complex is the essential part of the dimer linkage of genomic HIV-1 RNA. Proc Natl Acad Sci U S A 93, 5572–5577 (1996).

44. C. Wagner et al., Dimerization of the 3’UTR of bicoid mRNA involves a two-step mechanism. J Mol Biol 313, 511–524 (2001).

45. K. K. Alam, K. D. Tawiah, M. F. Lichte, D. Porciani, D. H. Burke, A Fluorescent Split Aptamer for Visualizing RNA-RNA Assembly In Vivo. ACS Synth Biol 6, 1710–1721 (2017).

46. P. Rangan et al., Temporal and spatial control of germ-plasm RNAs. Curr Biol 19, 72–77 (2009).

47. R. Lorenz et al., ViennaRNA Package 2.0. Algorithms Mol Biol 6, 26 (2011).

48. O. A. Shanaa, A. Rumyantsev, E. Sambuk, M. Padkina, In Vivo Production of RNA Aptamers and Nanoparticles: Problems and Prospects. Molecules 26, (2021).

49. M. Zubradt et al., DMS-MaPseq for genome-wide or targeted RNA structure probing in vivo. Nat Methods 14, 75–82 (2017).

50. S. Rouskin, M. Zubradt, S. Washietl, M. Kellis, J. S. Weissman, Genome-wide probing of RNA structure reveals active unfolding of mRNA structures in vivo. Nature 505, 701–705 (2014).

51. H. A. Curnutte et al., Proteins rather than mRNAs regulate nucleation and persistence of Oskar germ granules in Drosophila. Cell Rep 42, 112723 (2023).

52. K. E. Kistler et al., Phase transitioned nuclear Oskar promotes cell division of Drosophila primordial germ cells. Elife 7, (2018).

53. A. Dahanukar, R. P. Wharton, The Nanos gradient in Drosophila embryos is generated by translational regulation. Genes Dev 10, 2610–2620 (1996).

54. E. R. Gavis, R. Lehmann, Translational regulation of nanos by RNA localization. Nature 369, 315–318 (1994).

55. E. R. Gavis, L. Lunsford, S. E. Bergsten, R. Lehmann, A conserved 90 nucleotide element mediates translational repression of nanos RNA. Development 122, 2791–2800 (1996).

56. R. Chen, W. Stainier, J. Dufourt, M. Lagha, R. Lehmann, Repressor sequestration activates translation of germ granule localized mRNA. bioRxiv, 2023.2010.2017.562687 (2023).

57. S. Crucs, S. Chatterjee, E. R. Gavis, Overlapping but distinct RNA elements control repression and activation of nanos translation. Mol Cell 5, 457–467 (2000).

58. A. Dahanukar, J. A. Walker, R. P. Wharton, Smaug, a novel RNA-binding protein that operates a translational switch in Drosophila. Mol Cell 4, 209–218 (1999).

59. E. R. Gavis, D. Curtis, R. Lehmann, Identification of cis-acting sequences that control nanos RNA localization. Dev Biol 176, 36–50 (1996).

60. H. Eisenberg, G. Felsenfeld, Studies of the temperature-dependent conformation and phase separation of polyriboadenylic acid solutions at neutral pH. J Mol Biol 30, 17–37 (1967).

61. W. Ma, G. Zhen, W. Xie, C. Mayr, In vivo reconstitution finds multivalent RNA-RNA interactions as drivers of mesh-like condensates. Elife 10, (2021).

62. B. Van Treeck, R. Parker, Emerging Roles for Intermolecular RNA-RNA Interactions in RNP Assemblies. Cell 174, 791–802 (2018).

63. P. J. Tomezsko et al., Determination of RNA structural diversity and its role in HIV-1 RNA splicing. Nature 582, 438–442 (2020).

64. T. Trcek, H. Sato, R. H. Singer, L. E. Maquat, Temporal and spatial characterization of nonsense-mediated mRNA decay. Genes Dev 27, 541–551 (2013).

65. C. Wang, L. K. Dickinson, R. Lehmann, Genetics of nanos localization in Drosophila. Dev Dyn 199, 103–115 (1994).

66. N. Imamachi, K. A. Salam, Y. Suzuki, N. Akimitsu, A GC-rich sequence feature in the 3’ UTR directs UPF1-dependent mRNA decay in mammalian cells. Genome Res 27, 407–418 (2017).

67. D. Lavysh, G. Neu-Yilik, UPF1-Mediated RNA Decay-Danse Macabre in a Cloud. Biomolecules 10, (2020).

68. C. Wang, R. Lehmann, Nanos is the localized posterior determinant in Drosophila. Cell 66, 637–647 (1991).

69. R. Lehmann, C. Nusslein-Volhard, The maternal gene nanos has a central role in posterior pattern formation of the Drosophila embryo. Development 112, 679–691 (1991).

70. A. Dold et al., Makorin 1 controls embryonic patterning by alleviating Bruno1-mediated repression of oskar translation. PLoS Genet 16, e1008581 (2020).

71. E. R. Gavis, S. Chatterjee, N. R. Ford, L. J. Wolff, Dispensability of nanos mRNA localization for abdominal patterning but not for germ cell development. Mech Dev 125, 81–90 (2008).

72. K. M. Bhat, The posterior determinant gene nanos is required for the maintenance of the adult germline stem cells during Drosophila oogenesis. Genetics 151, 1479–1492 (1999).

73. A. Forbes, R. Lehmann, Nanos and Pumilio have critical roles in the development and function of Drosophila germline stem cells. Development 125, 679–690 (1998).

74. J. Ding et al., Visualizing RNA conformational and architectural heterogeneity in solution. Nat Commun 14, 714 (2023).

75. H. Maity, H. T. Nguyen, N. Hori, D. Thirumalai, Odd-even disparity in the population of slipped hairpins in RNA repeat sequences with implications for phase separation. Proc Natl Acad Sci U S A 120, e2301409120 (2023).

76. D. R. Micklem, J. Adams, S. Grunert, D. St Johnston, Distinct roles of two conserved Staufen domains in oskar mRNA localization and translation. EMBO J 19, 1366–1377 (2000).

77. D. Ferrandon, L. Elphick, C. Nusslein-Volhard, D. St Johnston, Staufen protein associates with the 3’UTR of bicoid mRNA to form particles that move in a microtubule-dependent manner. Cell 79, 1221–1232 (1994).

78. J. D. Laver et al., Genome-wide analysis of Staufen-associated mRNAs identifies secondary structures that confer target specificity. Nucleic Acids Res 41, 9438–9460 (2013).

79. S. Mohr et al., Opposing roles for Egalitarian and Staufen in transport, anchoring and localization of oskar mRNA in the Drosophila oocyte. PLoS Genet 17, e1009500 (2021).

80. R. E. Murphy, J. S. Saad, The Interplay between HIV-1 Gag Binding to the Plasma Membrane and Env Incorporation. Viruses 12, (2020).

81. S. M. Garcia, Y. Tabach, G. F. Lourenco, M. Armakola, G. Ruvkun, Identification of genes in toxicity pathways of trinucleotide-repeat RNA in C. elegans. Nat Struct Mol Biol 21, 712–720 (2014).

82. D. Musaev et al., UPF1 regulates mRNA stability by sensing poorly translated coding sequences. Cell Rep 43, 114074 (2024).

83. S. E. Bergsten, T. Huang, S. Chatterjee, E. R. Gavis, Recognition and long-range interactions of a minimal nanos RNA localization signal element. Development 128, 427–435 (2001).

84. D. M. Parker et al., mRNA localization is linked to translation regulation in the Caenorhabditis elegans germ lineage. Development 147, (2020).

85. A. H. Cardona et al., Self-demixing of mRNA copies buffers mRNA:mRNA and mRNA:regulator stoichiometries. Cell, (2023).

86. S. Adivarahan et al., Spatial Organization of Single mRNPs at Different Stages of the Gene Expression Pathway. Mol Cell 72, 727–738 e725 (2018).

87. A. Khong, R. Parker, mRNP architecture in translating and stress conditions reveals an ordered pathway of mRNP compaction. J Cell Biol 217, 4124–4140 (2018).

88. K. J. Venken et al., MiMIC: a highly versatile transposon insertion resource for engineering Drosophila melanogaster genes. Nat Methods 8, 737–743 (2011).

89. S. N. Manivannan et al., Targeted Integration of Single-Copy Transgenes in Drosophila melanogaster Tissue-Culture Cells Using Recombination-Mediated Cassette Exchange. Genetics 201, 1319–1328 (2015).

90. T. Trcek, T. Lionnet, H. Shroff, R. Lehmann, mRNA quantification using single-molecule FISH in Drosophila embryos. Nat Protoc 12, 1326–1348 (2017).

91. S. L. Dulk, S. Rouskin, Probing RNA Structure with Dimethyl Sulfate Mutational Profiling with Sequencing In Vitro and in Cells. J Vis Exp, (2022).

92. I. Gaspar, F. Wippich, A. Ephrussi, Enzymatic production of single-molecule FISH and RNA capture probes. RNA 23, 1582–1591 (2017).

93. P. Eastman et al., OpenMM 7: Rapid development of high performance algorithms for molecular dynamics. PLoS Comput Biol 13, e1005659 (2017).

94. J. D. Honeycutt, D. Thirumalai, The nature of folded states of globular proteins. Biopolymers 32, 695–709 (1992).

95. E. Marinari, G. Parisi, Simulated Tempering: A New Monte Carlo Scheme. Europhysics Letters (EPL) 19, 451–458 (1992).

96. M. R. Shirts, J. D. Chodera, Statistically optimal analysis of samples from multiple equilibrium states. J Chem Phys 129, 124105 (2008).

97. P. Debye, E. Hückel, Zur Theorie der Elektrolyte. I. Gefrierpunktserniedrigung und verwandte Erscheinungen. Physikalische Zeitschrift 24, (1923).

98. B. Lee, F. M. Richards, The interpretation of protein structures: estimation of static accessibility. J Mol Biol 55, 379–400 (1971).

99. S. Mitternacht, FreeSASA: An open source C library for solvent accessible surface area calculations. F1000Res 5, 189 (2016).

100. R. J. G. B. Campello, D. Moulavi, J. Sander, in Advances in Knowledge Discovery and Data Mining. (2013), chap. Chapter 14, pp. 160-172.

101. T. A. Bunch, Y. Grinblat, L. S. Goldstein, Characterization and use of the Drosophila metallothionein promoter in cultured Drosophila melanogaster cells. Nucleic Acids Res 16, 1043–1061 (1988).

102. K. Hanyu-Nakamura, K. Matsuda, S. M. Cohen, A. Nakamura, Pgc suppresses the zygotically acting RNA decay pathway to protect germ plasm RNAs in the Drosophila embryo. Development 146, (2019).

103. C. A. Schneider, W. S. Rasband, K. W. Eliceiri, NIH Image to ImageJ: 25 years of image analysis. Nat Methods 9, 671–675 (2012).

104. K. E. Deigan, T. W. Li, D. H. Mathews, K. M. Weeks, Accurate SHAPE-directed RNA structure determination. Proc Natl Acad Sci U S A 106, 97–102 (2009).

105. P. Kerpedjiev, S. Hammer, I. L. Hofacker, Forna (force-directed RNA): Simple and effective online RNA secondary structure diagrams. Bioinformatics 31, 3377–3379 (2015).

106. K. Darty, A. Denise, Y. Ponty, VARNA: Interactive drawing and editing of the RNA secondary structure. Bioinformatics 25, 1974-1975 (2009).

107. S. Bolte, F. P. Cordelieres, A guided tour into subcellular colocalization analysis in light microscopy. J Microsc 224, 213–232 (2006).

108. T. Lionnet et al., A transgenic mouse for in vivo detection of endogenous labeled mRNA. Nat Methods 8, 165–170 (2011).

109. M. de Cuevas, A. C. Spradling, Morphogenesis of the Drosophila fusome and its implications for oocyte specification. Development 125, 2781–2789 (1998).

